# StrainGE: A toolkit to track and characterize low-abundance strains in complex microbial communities

**DOI:** 10.1101/2021.02.14.431013

**Authors:** Lucas R. van Dijk, Bruce J. Walker, Timothy J. Straub, Colin J. Worby, Alexandra Grote, Henry L. Schreiber, Christine Anyansi, Amy J. Pickering, Scott J. Hultgren, Abigail L. Manson, Thomas Abeel, Ashlee M. Earl

**Affiliations:** Infectious Disease & Microbiome Program, Broad Institute, Cambridge, MA 02142, USA; Delft Bioinformatics Lab, Delft University of Technology, Van Mourik Broekmanweg 6, 2628 XE Delft, The Netherlands; Applied Invention, Cambridge, MA; Department of Immunology and Infectious Diseases, Harvard T.H. Chan School of Public Health, Boston, MA 02115, USA; Department of Molecular Microbiology, Washington University School of Medicine, St. Louis, MO 63110 USA; Center for Women’s Infectious Disease Research (CWIDR), Washington University School of Medicine, St. Louis, MO 63110 USA; Department of Civil and Environmental Engineering, University of California, Berkeley, Berkeley, CA, USA 94720; Stuart B. Levy Center for Integrated Management of Antimicrobial Resistance (Levy CIMAR), Tufts University, Boston, MA, USA

**Author notes:** Co-first author (Lucas R. van Dijk, Bruce J. Walker). Corresponding Author: Dr. Ashlee M. Earl, The Broad Institute, 415 Main Street, Cambridge, MA 02142, USA.

## Abstract

Human-associated microbial communities comprise not only complex mixtures of bacterial species, but also mixtures of conspecific strains, the implications of which are mostly unknown since strain level dynamics are underexplored due to the difficulties of studying them. We introduce the Strain Genome Explorer (StrainGE) toolkit, which deconvolves strain mixtures and characterizes component strains at the nucleotide level from short-read metagenomic sequencing with higher sensitivity and resolution than other tools. StrainGE is able to identify nearest known references and find variants for multiple conspecific strains within a sample at relative abundances below 0.1% in typical metagenomic datasets.

## Background

Human-associated microbial communities include complex mixtures of bacterial species. Many of these species are renowned for their genomic and phenotypic plasticity, which can lead to different human health outcomes. For example, strains of *Escherichia coli* can share less than 50% of their gene content and cause distinct disease including diarrhea and urinary tract infections, or potentiate tumorigenesis, while other strains are able to co-exist with their host without causing overt illness (1–3). Multiple distinct strains of the same species, often from across genetically dissimilar phylogroups, frequently coexist within a single human gut community (4,5), the implications of which are mostly underexplored due to the difficulties of studying strain-level variation from complex community samples.

While culture-based approaches have been a workhorse for dissecting strain-level diversity, these approaches can be slow and unfaithful to the true representation of strains, due to culturing bottlenecks that limit what diversity can be observed, as well as the potential for evolution during culture (6). Whole metagenome shotgun sequencing approaches offer less perturbed views of strain-level diversity, but require specialized tools to resolve. Current strain-level metagenomic data analytical tools (reviewed in Anyansi *et al.* (7)), including MIDAS (8) and StrainPhlan (9), have been successful in distinguishing strains present in different samples, but are unable to disentangle strain mixtures as they rely on a single nucleotide variant (SNV) profile per sample. Another class of tools, including BIB (10) and StrainEst (11), are designed to untangle strain mixtures, but are limited to reporting a close reference genome from a database, and are unable to distinguish between distinct strains that are close to the same reference. Tools including DESMAN (12) and inStrain (13) aim to recover strain-level variation after *de novo* metagenomic assembly, which is not always feasible for low abundance members of a community. To our knowledge, none of these tools alone can robustly disentangle mixtures of same-species strains, distinguish similar strains at the nucleotide level, and work robustly at the low coverages typically found for many clinically relevant organisms in metagenomic samples, as is typical for *E. coli* in the human gut (4).

We developed the Strain Genome Explorer (StrainGE) toolkit to disentangle strain mixtures, particularly when strains are present at low relative abundances, and to characterize and compare strains across samples at the nucleotide-level, with higher resolution than other tools. While what defines a strain depends on the research question to be answered (6), we created StrainGE to be sensitive, tunable and detailed in its output, enabling researchers to tailor it to their definition. We have extensively benchmarked StrainGE on synthetic data, compared StrainGE against other state-of-the-art strain detection tools, and applied StrainGE to multiple clinical human gut metagenomic datasets, demonstrating StrainGE’s ability to distinguish low-abundance strains in real metagenomic settings. Herein, we applied StrainGE to analysis of *E. coli* and *Enterococcus* strains, but StrainGE can be broadly applied to all community assemblages where same species bacterial strain dynamics are of interest.

## Results

### Strain Genome Explorer (StrainGE) toolkit

StrainGE is a toolkit for strain-level characterization and tracking of species (or genera) of interest from short read metagenomic datasets, tuned specifically to capture low abundance strains where data are scant. StrainGE has two key components: Strain Genome Search Tool (StrainGST), and Strain Genome Recovery (StrainGR). StrainGST sensitively reports reference genome(s) from a database that are most similar to the strain(s) in a sample. StrainGR analyzes short read alignments to a reported reference genome(s) to identify single nucleotide variants (SNVs) and large deletions (*i.e.,* gaps in coverage) relative to the reference. Though StrainGST can be used as a standalone tool, the StrainGE tool suite, including StrainGR, enables sensitive nucleotide-level comparison and tracking of strains across multiple samples and provides insights into potential functional variation among individual strains.

In brief, StrainGST builds a database of high-quality reference genomes (*e.g.,* RefSeq assemblies) from a species or genus of interest (Figure 1a), filtering them to remove highly similar genomes using a k-mer based clustering approach. To identify a similar reference(s) to the strain(s) within a sample and to estimate its relative abundance, StrainGST compares the k-mers in the sample to those of the database reference genomes (Figure 1b) using an iterative algorithm similar to that of QuantTB (14). StrainGR provides additional detailed information about gene content and SNVs in a strain(s) by analyzing alignments of metagenomic sequencing data to each StrainGST predicted reference (Figure 1c). To ensure accurate SNV calls while maintaining sensitivity at low coverage, StrainGR employs stringent quality thresholds and heuristics to filter spurious alignments and reduce the number of incorrect calls.

**Figure 1.**
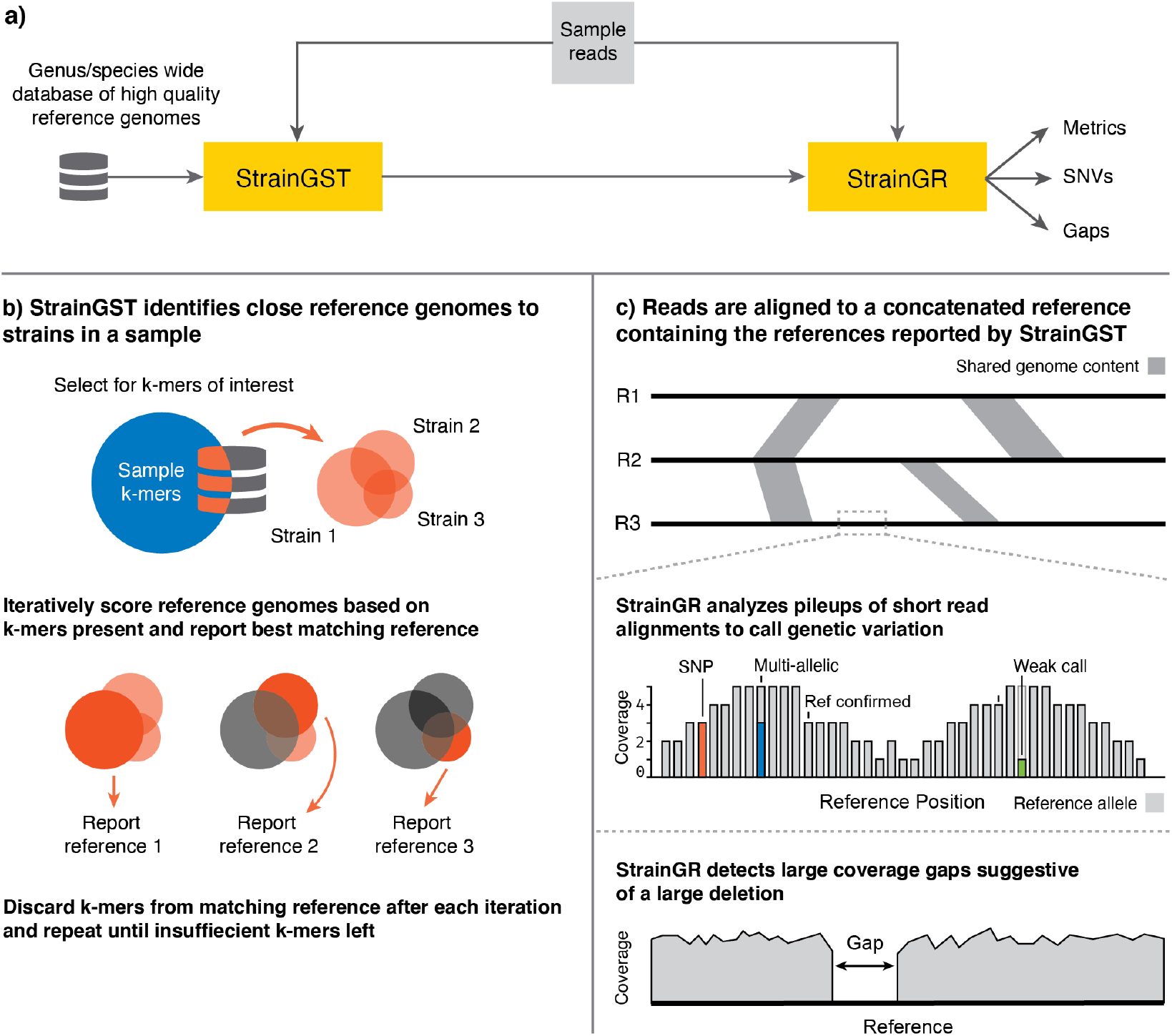
StrainGE is a toolkit to track, characterize and compare low-abundance strains in metagenomic samples. a) Overview of StrainGE pipeline. StrainGST uses a database of reference genomes to select those most similar to strains present in a metagenomic sample. StrainGR further characterizes SNVs and gaps that differ between references selected by StrainGST and the actual strain present in the sample. b) At each iteration, StrainGST scores each reference strain by comparing the k-mer profile of the reference to the sample k-mers, reporting the reference closest to the highest abundant strain in the sample. The k-mers in the reported reference are removed from the sample and the process is repeated to search for lower-abundance strains, until there are insufficient k-mers. c) StrainGR uses a short read alignment-based approach to characterize variation (SNVs and gaps) between the reference(s) identified by StrainGST and the metagenomic sample. Regions shared between the concatenated genomes (grey shaded areas) are detected and excluded from variant calling. After applying rigorous QC, positions in the reference are classified as i) “reference confirmed” (light grey; a single strong allele which matches the reference); ii) “SNV” (red; a single strong allele that is different than the reference); or iii) “multi-allelic” (blue; multiple strong alleles present, e.g. the blue allele together with the reference allele in grey). The position with a strong reference allele and a weak allele (green; an allele with only limited support in the reads) is classified as “reference confirmed” because only the reference allele is considered strong at that position.

To separate SNVs belonging to different strains, StrainGR creates a concatenated set of reference genomes, containing all references predicted by StrainGST in a sample or set of samples. It uses this reference set to align metagenomic reads and call variants. To prevent assigning alleles incorrectly, StrainGR only calls variants in regions unique to a single reference by filtering out ambiguously aligned reads. In cases where StrainGST has identified distinct but closely related strains across samples, StrainGR can perform another, coarser round of reference clustering prior to concatenation in order to increase the amount of unique sequence for variant calling.

These variant calls can in turn be used to compare strains across samples. StrainGR compares positions in the combined reference where both samples have a “strong” allele call (the common callable genome), roughly equivalent to positions where both samples have at least two good reads and >10% of the pileup supporting that allele. Strain relationships can be assessed using two key metrics: i) the Average Callable Nucleotide Identity (ACNI), or the percentage of common callable positions where both samples have a single identical base call; and ii) a “gap similarity” metric, as patterns of large deletions are often conserved between closely related strains, which can provide an orthogonal metric of strain similarity (15). Bringing StrainGST and StrainGR together, StrainGE can identify, track and characterize strains, including strain mixtures, at coverages as low as 0.5x across the genome, or approximately 0.08% relative abundance for a 5 Mb genome in 3 Gb of raw sequencing data.

#### *Benchmarking StrainGE on* Escherichia

StrainGE was designed to be broadly applicable across different bacterial genera and species, including less well-studied species lacking numerous high quality reference genomes (Supplemental Results A). For benchmarking, we focused on *E. coli*, an evolutionarily and functionally diverse species composed of eight main phylogroups (A, B1, B2, C, D, E, F, G; with mean within-phylogroup average nucleotide identity (ANI) ranging from 97.6%-99.1%) (16). Despite their importance to human health as opportunistic pathogens and frequent commensals, *E. coli* are typically found at low (<1%) relative abundance in diverse strain mixtures in human guts (4). Thus, *E. coli* presents a challenging and clinically relevant case on which to benchmark StrainGE.

We first used StrainGST to construct an *Escherichia-specific* reference database by downloading all available complete *Escherichia* assemblies from NCBI RefSeq (929 at time of download; Materials and Methods; Supplemental Table 1). Because we were interested in tracking specific strains as opposed to plasmids, and because plasmids readily transfer between different genetic backgrounds of the same and/or different species (see Supplementary Results A) (17), scaffolds labeled as plasmid, or those smaller than 1 Mbp were removed. To select unique database representatives, StrainGST uses a k-mer based clustering method to cluster genomes with ANI higher than approximately 99.8% (a tunable parameter; optimization in Supplemental Table 4) to other reference(s) in the database. The resulting StrainGST database contained 361 complete *Escherichia* chromosomes, comprising 341 *E. coli* and *Shigella* chromosomes, representing all eight phylogroups, and 20 chromosomes from other *Escherichia* species (Supplemental Table 1).

#### StrainGST works at lower coverages and pinpoints more closely related references than other tools

In order to assess the sensitivity and specificity of StrainGST compared to similar tools like BIB (10) and StrainEst (11), we constructed *in silico* metagenomes that were spiked with sequences of known strains of *Escherichia* at varying relative abundances. StrainGST performed as well as, or better than, the other tools across all scenarios tested, and stood out strongly when strains were at very low abundance, either alone or as part of a mixture with other strains (Supplementary Results B). StrainGST had the highest precision (mean 0.99) and F1 score (mean 0.99), and its recall and average Mash similarity (18) were at least as good as that of the other tools, when given the task of identifying the closest reference(s) to those present in the spiked metagenome (Supplementary Figure 1).

To assess the performance of each tool on a real sample of known composition, we created and sequenced a mock community containing approximately 99% human DNA and 1% *E. coli* DNA, representing a mixture of four distinct, previously sequenced strains with fully finished genomes mixed in unequal (80:15:4.9:0.1) relative abundances (Materials & Methods). Two of these strains were represented in the reference database, while the other two had approximate ANI similarities (or 1 - Mash distances (18)) of 99.95% and 99.89% to their closest representative in the database. StrainGST was the only tool that correctly identified all four *E. coli* strains, with no additional predictions (Table 1). While StrainEst correctly identified the same four strains, it also reported two false positive strains not present in the mixture. BIB’s limited database lacked references as closely related as the other two tools, but it was able to identify the closest matching reference in its own database; however, it also reported six false positive strains.

**Table 1.** StrainGST was the only tool that correctly identified the mock community composition.

| Predicted Strains (approximate ANI to true strain)* |  |  |  |
| --- | --- | --- | --- |
| Actual Strain present in mock community (phylogroup) | StrainGST | StrainEst | BIB |
| *E. coli* SEC470 (A) | ✓ | ✓ | *E. coli* K-12 GM4792, 99.24 % |
| *E. coli* UTI89 (B2) | *E. coli* UM146; 99.95% | *E. coli* UM146, 99.95% | *E. coli* H105, 98.49% |
| *E. coli* Sakai (E) | *E. coli* 149, 99.89% | *E. coli* 149, 99.89% | *E. coli* 108, 99.97% |
| *E. coli* 24377A (B1) | ✓ | ✓ | *E. coli* S40, 99.01% |
|  |  | ✗ *E. coli* APEC IMT5155 | ✗ *S. flexneri* G1663 |
|  |  | ✗ *E. coli* RM14721 | ✗ *E. coli* LHM10-1 |
|  |  |  | ✗ *E. coli* MSHS 133 |
|  |  |  | ✗ *S. dysenteriae* 80-547 |
|  |  |  | ✗ *E. coli* IMT16316 |
|  |  |  | ✗ *S. dysenteriae* ATCC 12039 |

\*A check mark indicates that the exact strain was present in our database and correctly identified. A strain name on a green background indicates that the exact reference was not in the reference database, but the closest available reference was correctly identified (along with its approximate ANI to the actual strain). An “X” on a yellow background indicates a false positive strain identified by the tool that was not present in the mock community.

StrainGST predicted that *E. coli* represented 0.74% of the total sample, which was close to the target value (1%) and consistent with predictions from MetaPhlAn2 (19) and Bracken (20). StrainGE also reported predicted abundances of the four detected strains, which were in the expected range. The capability to predict relative abundances was unique to StrainGE and not available for the other tools.

#### StrainGE was the most accurate at detecting shared strains at coverages as low as 0.1x

StrainGST uses similarity to reference genomes as a proxy for strain identity and to calculate relative abundances of strains in a sample. However, the success of this approach to identify the “same” versus “different” strain varies with the granularity of the reference database. That is, StrainGST with a larger reference database will be more specific to detect distinct strains than with a smaller reference database. StrainGR was designed to complement StrainGST by providing a more detailed view of the nucleotide- and gene-level differences between a strain in a sample and its closest reference, which can be used to compare across samples having strains that match the same reference. While a close reference genome generally results in more accurate alignments and variant calls (21), StrainGR still provides meaningful relationships when the reference is more distant, as would be the case in a smaller constructed database or with less well-studied organisms (Supplemental Results A). Thus, StrainGR extends the tool’s sensitivity to distinguish different strains and minimizes biases introduced in database construction.

Using *Escherichia* spiked metagenomes, we first benchmarked the ability of StrainGR to call variants—*i.e.,* SNVs and deletions (or gaps in coverage)—against close references for *Escherichia* strains present in each sample. StrainGE accurately called SNVs and large deletions, for both single strain and mixture samples, providing key information to assess whether two samples share a strain (Supplementary Figure 2; Supplementary Results C-D).

Next, we selected two recent and highly cited strain-tracking tools, MIDAS (8) and StrainPhlan (9), to compare with StrainGR’s performance in tracking identical strains across samples containing one or more strains. To assess the sensitivity of these tools to distinguish between similar strains, we generated pairs of spiked metagenomic samples, each containing one or more *Escherichia* strains at between 0.1x and 10x coverage. Similar strain pairs were derived from the same reference genome, but with a different set of ~5,000 random SNVs introduced *in silico* into each strain’s genome (Figure 2a-b). This resulted in each strain having 99.9% ANI to the reference and each strain pair having 99.8% ANI to one another. This identity level should result in strain pairs matching the same StrainGST reference but still distinguishable by StrainGR. By default, MIDAS and StrainPhlan require that a strain have at least 5x and 10x coverage, respectively, to run to completion, which is higher than that required by StrainGE. Thus, we ran each of these tools using default settings, in order to evaluate their default performance at higher coverages, as well as with manual tuning to allow the tools to accommodate our lower coverage benchmarks (Materials & Methods).

**Figure 2.**
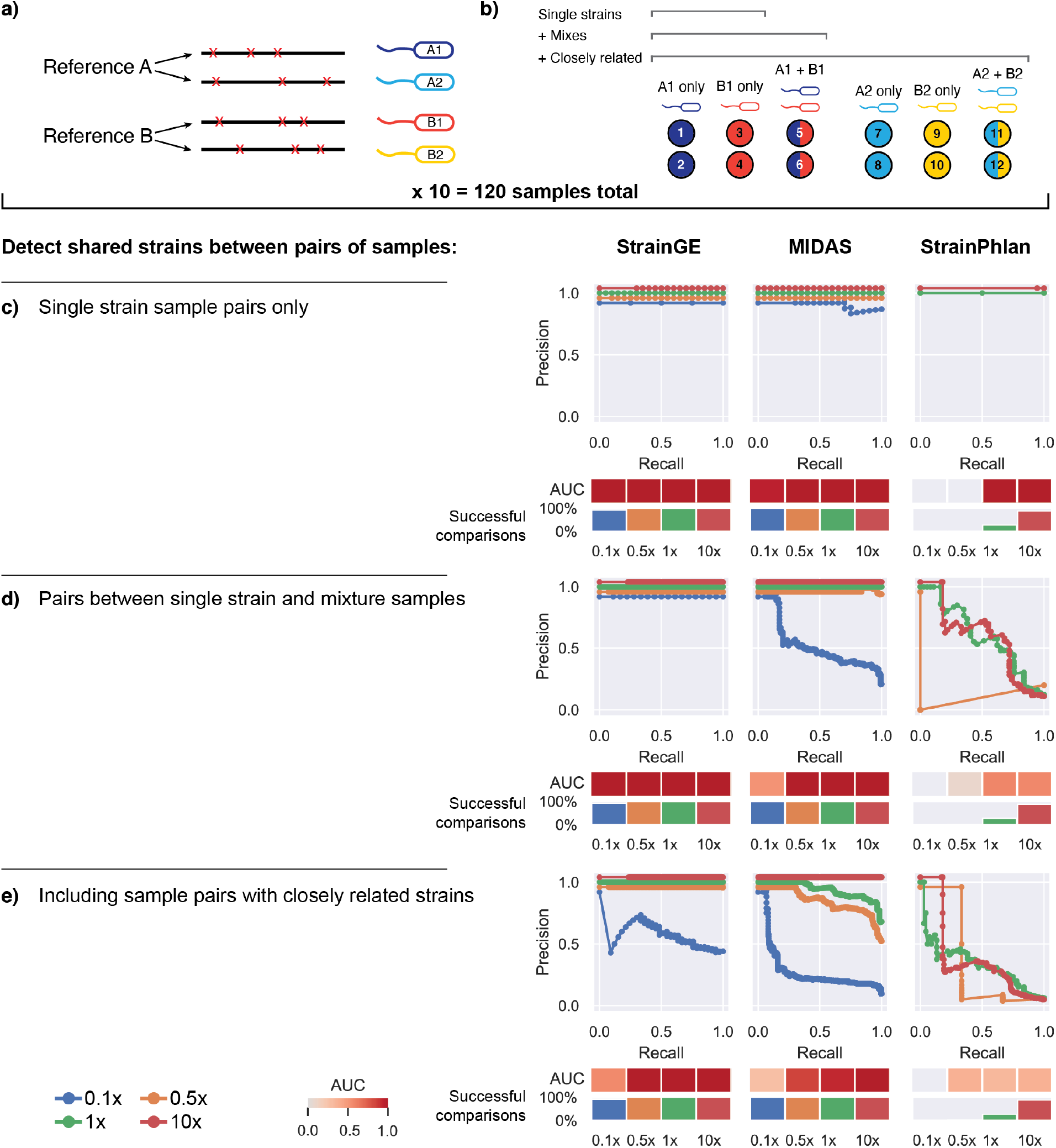
StrainGE is the only tool that can detect strain sharing at coverages as low as 0.1x. a) Depiction of how synthetic *Escherichia* genomes were generated from randomly selected NCBI RefSeq genomes to create sets of closely related strains (e.g., A1/A2 and B1/B2) for spike in experiments. b) Depiction of how spiked metagenomes were created using synthetic genomes from (a). Each circle represents a spiked metagenome. The color of the circle indicates which synthetic strain was included: single color circles indicate spiked metagenomes containing a single synthetic strain, and two color circles indicate spiked metagenomes containing two synthetic strains mixed at equal proportions. c-e) Precision-recall curves for each tool and coverage 0.1x-10x, when given the task to detect which sample pairs contain identical strains. The area under the curve (AUC) is depicted as a heatmap below. The “successful comparisons” bar plot indicates the percentage of sample pairs for which a comparison was possible (*i.e.*, tools ran to completion for both samples). c) Limiting to single-strain samples from distinct references. d) Including samples with two strains, but limited to strains from distinct references. e) Including samples with closely related strains.

At 10x coverage, MIDAS and StrainPhlan performed comparably using default and tuned settings (Figure 2; Supplemental Figure 3). While StrainGE and MIDAS performed well across all scenarios at high coverage, StrainPhlan performed poorly on mixes because it only reported a single SNV profile for each sample. For lower coverage scenarios, StrainGE consistently outperformed the manually tuned versions of MIDAS and StrainPhlan (Figure 2). For single strain samples, StrainGE perfectly matched strain pairs down to 0.1x coverage, with MIDAS performing comparably (Figure 2c). StrainPhlan performed marginally at 1x coverage, and still was unable to run to completion at coverages lower than 1x. For simple mixtures (Figure 2d), only StrainGE and MIDAS correctly matched most pairs, because StrainPhlan was unable to disentangle mixes. StrainGE was the only tool that was able to generate results across the whole range of coverages, scoring almost perfectly even down to the lowest tested coverage of 0.1x. When we included samples containing very closely related pairs (Figure 2e), StrainGE and MIDAS performed well down to 0.5x coverage, but StrainPhlan could not distinguish between closely related strains, even at 10x coverage, likely due to its reliance on marker genes which comprise only a small fraction of the genome.

Aside from its relative high sensitivity, another key advantage of StrainGE over the other tools is its ability to link a strain in a sample to its specific close reference genome reported by StrainGST, which places an observed strain within the known phylogenetic structure of the reference database (Supplemental Figure 4). In contrast, the SNV profiles output by StrainPhlan (based on marker genes) or MIDAS (compared to a single built in *E. coli* reference), do not offer convenient phylogenetic placement.

#### StrainGE can be used to approximate the ANI between low-abundance strains within metagenomic samples

Having demonstrated that StrainGE is able to track strains across samples, we next sought to determine how well StrainGE metrics recapitulated genetic relationships between strains, as well as to evaluate how other *Escherichia* strains in a sample might influence these metrics. We generated another set of spiked metagenomes similar to that described above, except we spiked into each metagenome one of a set of diverse known *E. coli* genomes with between 0 and 5,000 random SNVs introduced *in silico* (100%-99.9% ANI to reference, steps of 0.01%). Individual strains were spiked into multiple metagenomic samples (potentially containing other *E. coli* strains) at random coverage levels of 0.1x-10x. We then ran StrainGE on each sample, and compared strains that matched the same close reference via ACNI, computed from across the common callable regions of the reference, and gap similarity scores.

Identical strains in different samples were found to have high ACNI and gap similarity (Figure 3a), despite different metagenomic backgrounds. StrainGR’s ability to correctly delineate identical strain pairs increased with a larger common callable genome (Supplementary Figure 5), with a substantial drop in accuracy when the common callable genome fell below 0.5%. Of the same-strain pairs with at least 0.5% common callable genome, 74% had an ACNI >99.98% and 96% had a gap similarity >0.98, highlighting StrainGE’s ability to identify same-strain pairs even at low abundance and in the presence of other strains.

**Figure 3.**
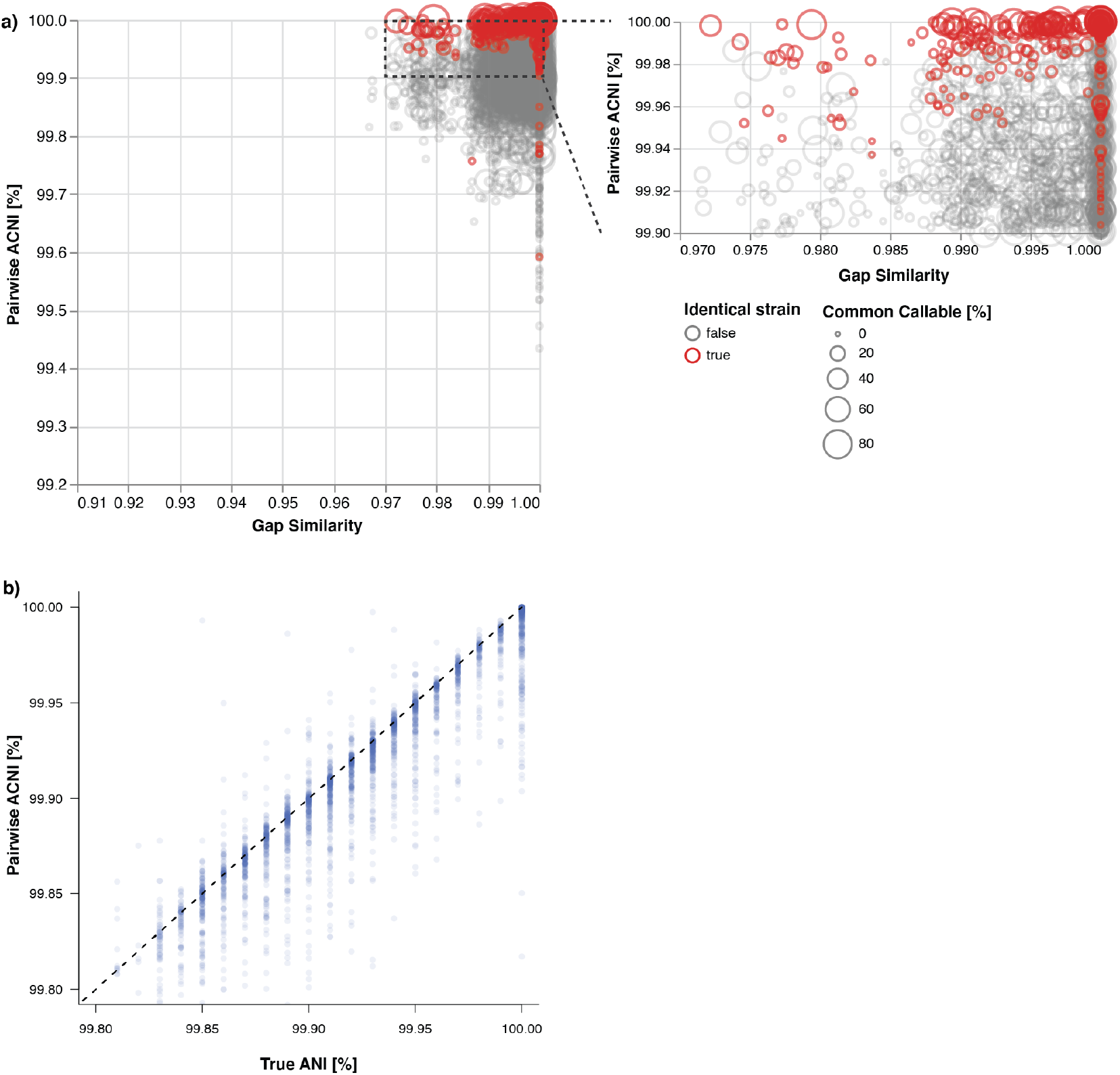
StrainGR can discriminate between highly similar strains and reports ACNI values that correlate well with the true ANI between strains. a) For all synthetic sample pairs with the same StrainGST reference called, the Jaccard gap similarity index and pairwise ACNI are plotted. Circle size indicates the percentage of the reference genome that was callable across both strains being compared. Red circles indicate comparisons between identical strains. b) For all pairs, the true ANI between spiked isolates is plotted against the ACNI, as estimated by StrainGR. The dashed line indicates parity between these metrics.

StrainGR’s ACNI measurements across all strain pairs (that can have 0-10,000 SNV differences) also correlated strongly with true ANI, even though ACNI is based on unique regions, and ANI is based on the entire genome. Ninety two percent of ACNI values were within 0.05% of the true ANI, regardless of metagenomic background though ACNI values in this benchmark tended to fall below the true ANI value (Figure 3b).

### In real metagenomic data, StrainGE identifies low abundance strains and can track strains across samples, including in strain mixtures

#### StrainGE can identify lower abundance instances of persistent strains previously undetectable by other tools

In order to assess StrainGE’s utility to characterize strains from real world samples, we examined its performance, using default parameters with our *Escherichia* reference database, on a previously published metagenomic dataset of 27 longitudinally collected stool samples from a patient with Crohn’s disease, upon which MIDAS was run to delineate *E. coli* strains (22). MIDAS identified seven dominant “strain types” (“ST1” - ”ST7”) that varied in abundance over time. Each of these belonged to a distinct multi-locus sequence type (MLST) and represented one of five *E. coli* phylogroups. StrainGE showed good concordance with results from MIDAS for all of the high abundance strains (>10% abundance) (Table 2). For the two calls that disagreed, our StrainGST database lacked representatives for the two MLSTs reported by MIDAS. However, StrainGE selected the next closest reference, which we confirmed was correct by comparing the whole genome sequence from a cultured representative of ST1 reported in Fang *et al.* to our reference database.

**Table 2.** The strains predicted by MIDAS match the dominant strains predicted by StrainGE.

| Strain<br>(time points) | MIDAS<sup>§</sup> |  | StrainGST |  |  |
| --- | --- | --- | --- | --- | --- |
|  | MLST | *E. coli*<br>phylogroup | MLST | *E. coli*<br>phylogroup | Most abundant strain |
| **ST1** (1) | 95 | B2 | 95 | B2 | PA45B* |
| **ST2** (2) | 1629<sup>†</sup> | E | 1011 | E | Santai |
| **ST3** (3,4) | 69 | D | 69 | D | 118UI |
| **ST4** (11-18) | 58 | B1 | 58 | B1 | D5 |
| **ST5** (19-22,<br>27) | 131 | B2 | 131 | B2 | MVAST0167 |
| **ST6** (23,24) | 409<sup>†</sup> | A | 1408 | A | AR_0061 |
| **ST7** (25,26) | 1727 | B1 | 1727 | B1 | 2011C-3911 |

§ Fang *et al.* (22)
\* The actual strain corresponding to ST1 (3\_2\_53FAA) was whole-genome sequenced by Fang *et al.* PA45B and 3\_2\_53FAA share 99.9% average nucleotide identity based on whole-genome comparative genomics analysis.
† MLST profile was not represented by any reference genome in the StrainGE database. StrainGST predicted the closest reference within the StrainGE database, which was within the same phylogroup.

StrainGST also identified seven distinct strains that were missed by MIDAS (Figure 4a). While the majority of these were secondary strains found to coexist with a dominant strain predicted by MIDAS, StrainGST also predicted strains at timepoints where MIDAS called none (time points 6-10). In most of these cases, the strains were at or below 1% relative abundance and were also detected by MIDAS at higher abundance in other time points (*e.g.,* ST3; dark green), lending credence to their existence in these samples and suggesting that some strains were more persistent over time than previously predicted (Figure 4a).

**Figure 4.**
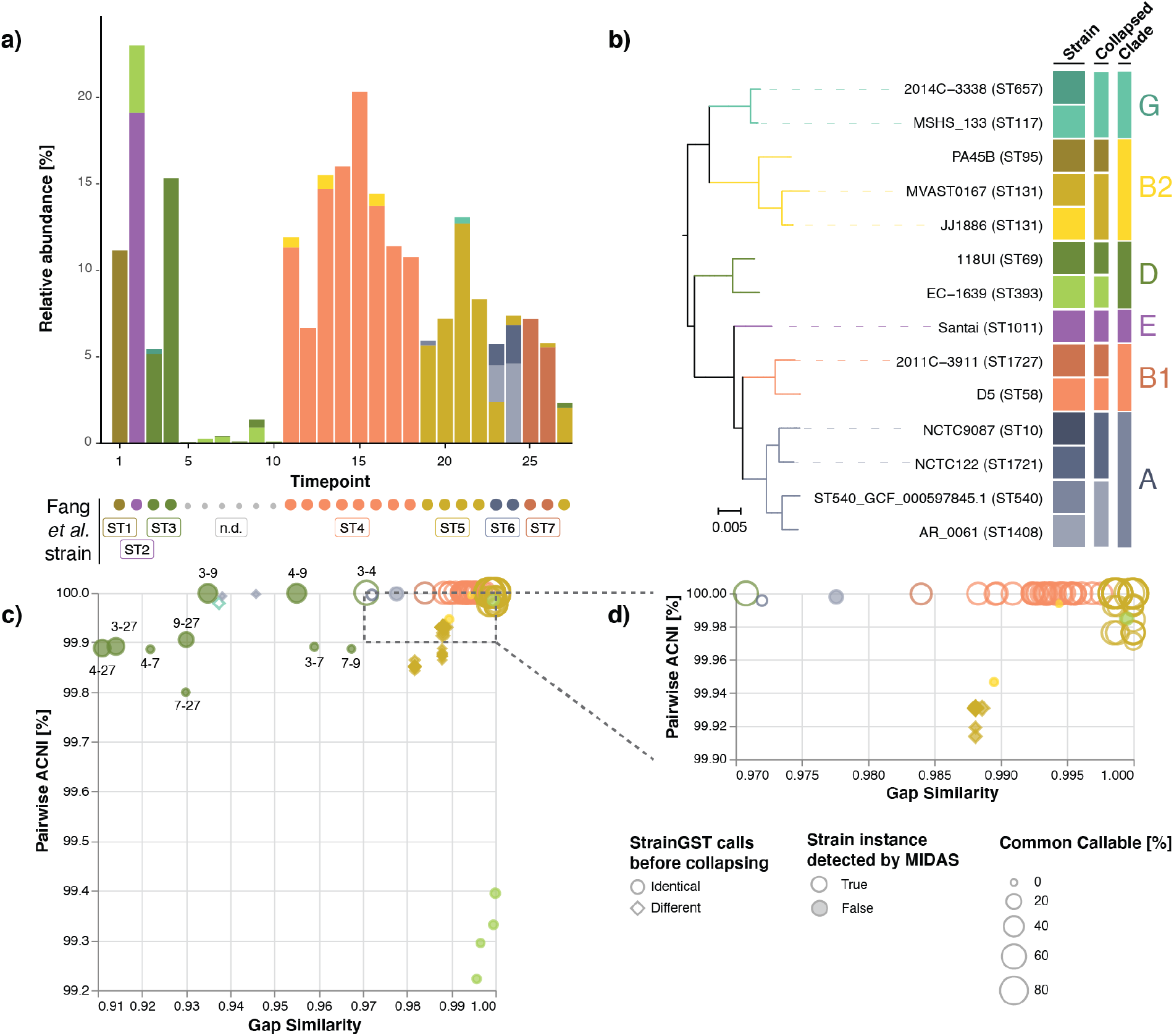
StrainGE accurately and sensitively identifies distinct *E. coli* strains in a longitudinal collection of stool sample metagenomes from an individual with Crohn’s Disease. a) Stacked barplot showing the relative abundances of StrainGST calls for each of 27 longitudinal stool metagenomes from Fang *et al.* Circles indicate the strain detected in Fang *et al.*, colored by its StrainGST counterpart and labeled using the ST designations (ST1-ST7) assigned by Fang *et al.* Small grey circles indicate samples where no strain was predicted in Fang *et al.;* these are labeled with “n.d.” b) Single-copy core phylogeny of the 14 StrainGST reference genomes with close matches to strains across samples. “Strain” colors are based on the clade from which each strain is predicted to belong; see column “Clade”. “Collapsed” column indicates which reference was selected as a representative for subsequent StrainGR analysis, when two or more references shared more than ~99.2% ANI. c) For all sample pairs matching the same collapsed reference, the Jaccard gap similarity index and pairwise ACNI are plotted. Circles indicate comparisons where the predicted reference was the same before collapsing, and diamonds indicate cases where the predicted reference before collapsing was different. Sizes of shapes indicate the percentage of the reference genome that was callable across both strains being compared. Filled in shapes indicate whether this strain instance was undetected by MIDAS. Dark green circles are labelled with the timepoints compared. d) Zoomed in view of the upper right corner of figure c).

After collapsing references to include a single representative from each closely related group of references (approximately 99.2% or greater ANI; ensuring every genome had at least 20% unique content) (Figure 4b; phylogroups G, B2 and A), StrainGR was run on all datasets using a concatenated reference including 10 references, representing the 14 strains detected by StrainGST. SNV and gap patterns predicted by StrainGR showed that the majority of strains matching the same reference had strikingly high pairwise ACNI (>99.96%) and gap similarity (>0.97) (Figure 4c,d), which were within the range of those of same-strain sample pairs in our simulations (Figure 3a). However, StrainGR results from strains matching the *E. coli* 118UI (dark green) reference stood out. While 118UI-like strains from samples 3 and 4 had ACNI and gap similarity relationships that were on par with what we observed in same-strain simulations, all other comparisons fell outside of this range, suggesting that this individual carried a mixture of 118UI-like strains in their gut over time that were closely related, but not necessarily the same with respect to gene content and single nucleotide variation (Supplementary Figure 6).

#### *StrainGE accurately and sensitively identified a low-abundance, persistent strain of* E. coli *in longitudinal stool samples from a woman with recurrent urinary tract infection*

Although the results of StrainGE on the Fang *et al.* (22) dataset highlighted its ability to resolve strains present at low abundance, the overall *E. coli* relative abundances in these samples were significantly higher (median 7.9%; range 0.05%-27%) than those typically seen in the human gut. Thus, we also tested StrainGE on 12 stool metagenomes having more typical *E. coli* relative abundances (median 0.55%; range 0.006%-17.4%), which originated from a single individual with a history of recurrent urinary tract infection (rUTI) over the span of a year. Given that the gut is a known important reservoir for UTI-causing *E. coli* (23), it was of interest to trace gut *E. coli* strain dynamics and their relationship with UTI.

StrainGST detected a total of five distinct strains of *E. coli* (Figure 5a), including a recurrent strain detected in over half of samples. The persistent strain, an *E. coli* 1190-like strain from phylogroup D, had a median relative abundance of only 0.6% (range 0-1.2%) and was detected even in samples that were composed of multiple *E. coli* strains, including at very low (20-fold less) abundance relative to another strain (Figure 5a). Despite its low abundance, we were able to confirm that all *E. coli* 1190-like strains had extremely high ACNI (>99.95%) and gap similarity (>0.98) (Figure 5b), in line with the identities observed for same-strain benchmarking (Figure 3a), suggesting that this strain, also the causative agent of this individual’s rUTI, persisted long-term in their gut.

**Figure 5.**
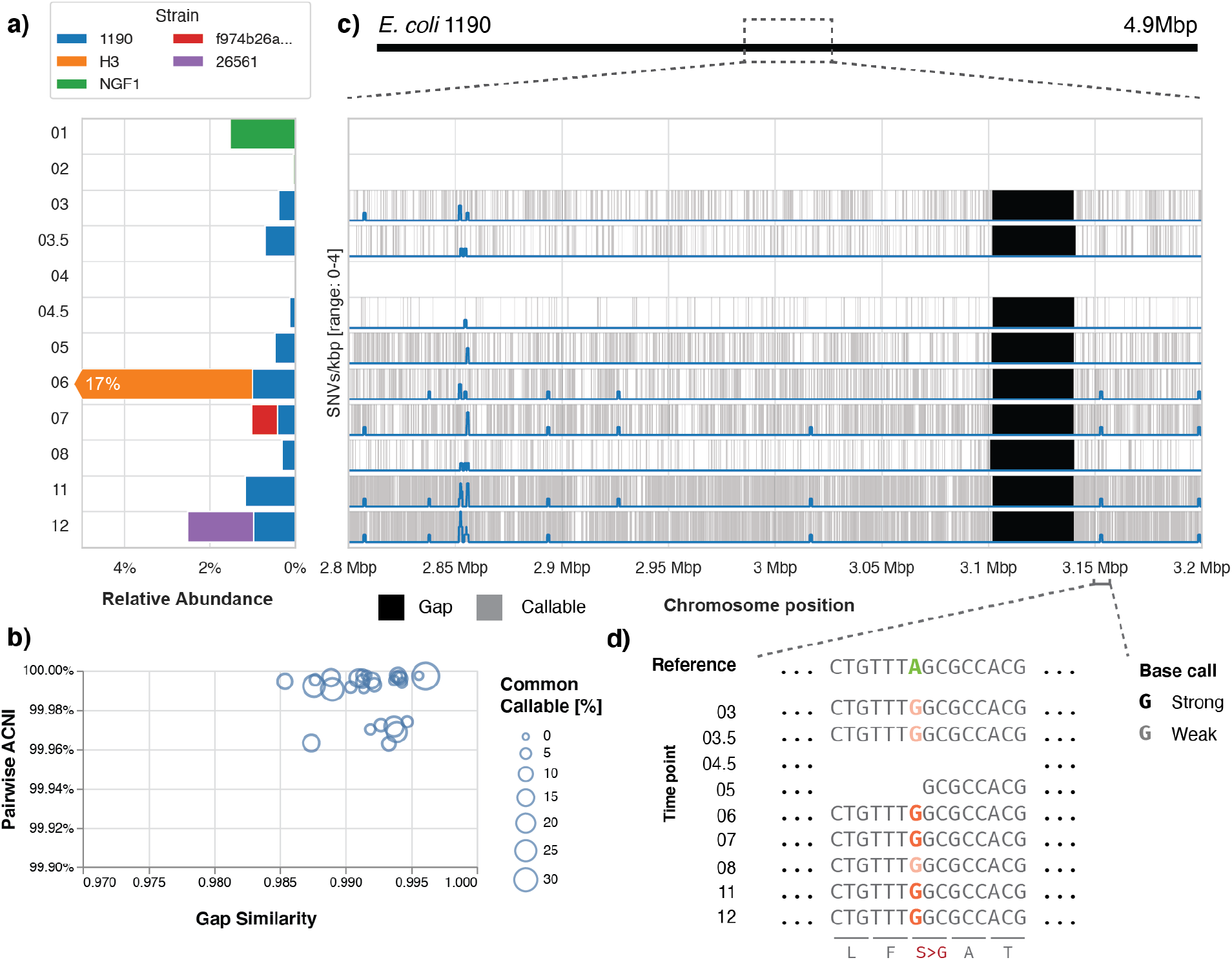
StrainGE detected a long-term, persistent strain of *E. coli* in a woman with rUTI. a) Relative abundances predicted by StrainGE are shown for all *E. coli* strains detected. b) For all sample pairs containing a strain matching to *E. coli* 1190, plot shows pairwise ACNI and gap similarity scores. Size of the circle indicates the percentage of the common callable genome. c) Zoom in of a region of the chromosome of *E. coli* 1190. Grey shaded areas indicate “callable” regions, where StrainGR had enough read data to make a strong allele call. Predicted gaps are shaded black. The blue line represents the number of SNVs per 1,000 bp, observed in at least 3 samples. d) Further zoom-in representing a region where StrainGR identified a nonsynonymous SNV that was consistently detected across all 1190-like strains.

Further, StrainGR output enabled us to look closely at the locations and identities of SNVs and genes within gaps relative to the reference. For example, we consistently identified a large gap across all time points encoding a prophage found in the original reference, but apparently lacking in the *E. coli* 1190-like strain in this individual (Figure 5c). Using StrainGR output that included both strong and weak variant calls (see Figure 1c for strong vs. weak calls; Materials & Methods), we were able to track 839 variant sites across samples, where the corresponding allele was strongly called in at least one sample, and weakly called in at least five samples (*e.g.,* a nonsynonymous SNV in the gene *cydC*; Figure 5d). At each of the 839 variant sites, the called allele was identical across all time points, except for three sites where another secondary weak allele was called, further supporting the persistence of a single UTI-causing strain.

#### StrainGE accurately recapitulated known strain level diversity and traced strains from mother to child

To demonstrate StrainGE’s applicability to other bacterial genera, we selected a previously published dataset investigating the impact of mode of delivery on the infant gut microbiome, including transmission and carriage of opportunistic pathogens from the *Enterococcus* genus (24). Shao *et al.* longitudinally followed 596 babies (and 175 mothers) by collecting stool samples that were then whole metagenome shotgun sequenced and cultured for pathogens, including 451 enterococci that were then whole genome sequenced. This dataset allowed us to evaluate StrainGE’s ability to report on i) the relationships between enterococcal strains predicted directly from metagenomes in comparison to those calculated from the genomes of cultured isolates, and ii) mother and child strain sharing. Further, in comparison to *Escherichia*, there were relatively few RefSeq complete *Enterococcus* genomes (283 *Enterococcus* vs. 929 *Escherichia*) with the numbers dwindling substantially for any given species *e.g.*, *Enterococcus faecalis* was represented by only 54 genomes (in comparison to 911 for *E. coli*). Thus, this dataset also allowed us to evaluate StrainGE’s ability to predict and compare strains across samples using a less dense database.

Using the same parameters as for the *Escherichia* database, we built a 163-member StrainGST database representing references from 80 *E. faecium*, 39 *E. faecalis* and 44 other enterococcal species (Supplemental Table 1). Next, we downloaded and ran StrainGST on all 1,679 stool metagenomes. StrainGST reported 2,024 instances of strains matching to 107 references, with the species distribution of StrainGST results roughly similar to the species distribution of the sequenced enterococcal isolates, as reported by Shao *et al.* (Table 3).

**Table 3.** Species distribution of reported StrainGST references and bacterial isolates.

| Species | StrainGST | Isolates |
| --- | --- | --- |
| *E. faecalis* | 64.5% | 78.7% |
| *E. faecium* | 8.4% | 13.7% |
| *E. avium* | 6.9% | 0% |
| *E. casseliflavus* | 6.6% | 2.3% |
| *E. durans* | 3.9% | 3.6% |
| Other enterococci | 9.7% | 1.7% |

One striking finding from this published study was the frequent isolation of closely related enterococci belonging to five *E. faecalis* lineages (ST16, ST30, ST191, ST40 and ST179) across different babies and hospitals (Figure 6a). To examine how well StrainGE could recapitulate these strain relationships, we ran StrainGR on each metagenome, using a concatenated reference including all StrainGST reported references per subject (and including the mother, if available), to determine ACNI and gap similarity for sets of strains that matched the same reference. Our results echoed the original finding of many closely related strains from the same lineages; 50% of *E. faecalis* strain comparisons had an ACNI >99.95% (Figure 6b). Further, nearly half (42%) of StrainGST strain predictions belonged to one of the five previously reported *E. faecalis* lineages, though our reference database lacked a ST179 representative and reported the closest match, ST711, instead.

**Figure 6.**
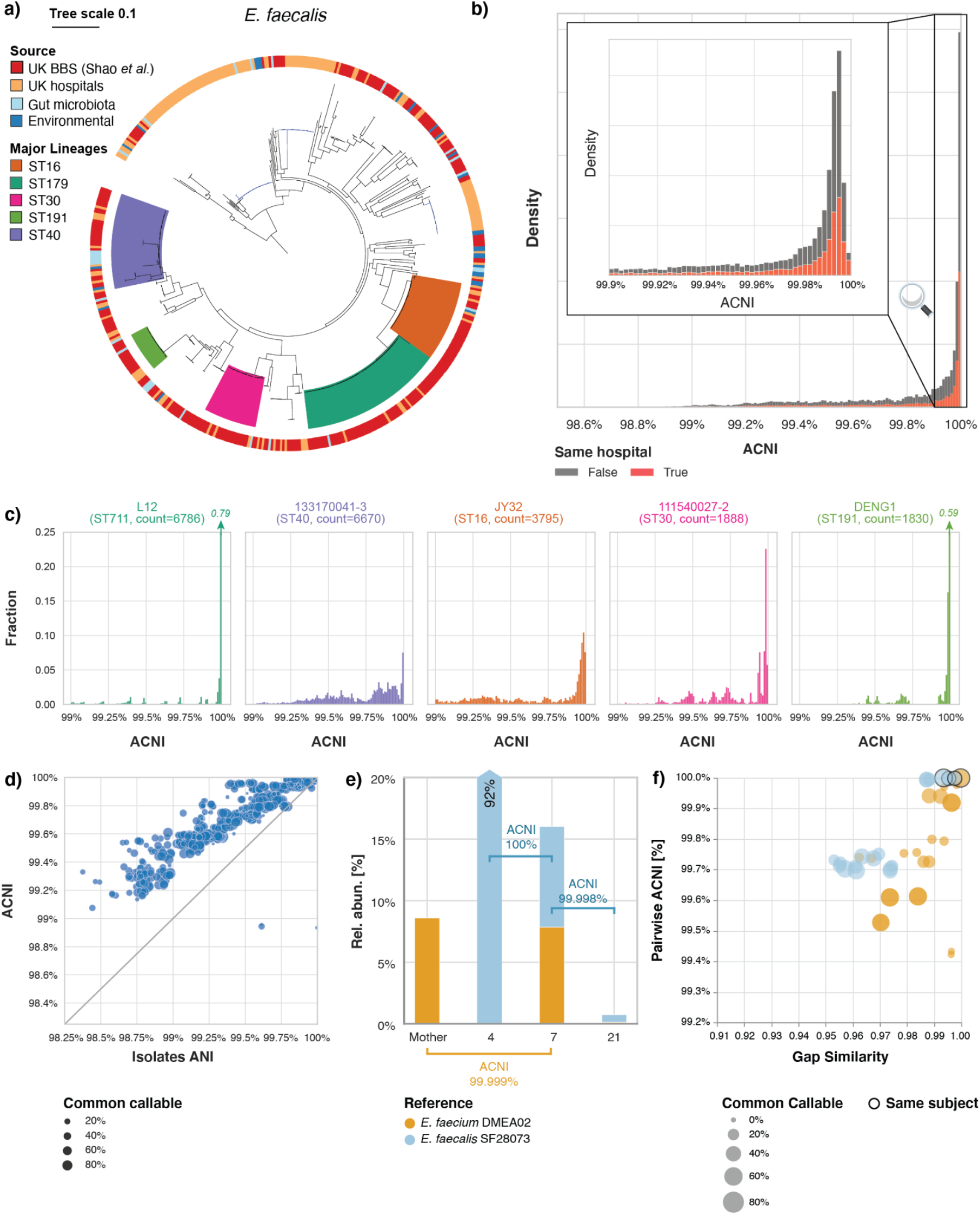
StrainGE recapitulates strain-diversity among bacterial isolates using metagenomic data only. a) Single-copy core phylogenetic tree of *E. faecalis* isolates from the UK Baby Biome Study (UK BBS) (*n*=282) in the context of isolates from other public UK hospitals (*n*=168), human gut microbiota (*n*=28), or other environmental sources (*n*=27). Five major lineages were identified, represented by ST16, ST179, ST30, ST191 and ST40. Tree republished with permission from Shao *et al.* (24). b) Pairwise ACNI distribution of all pairwise strain comparisons, colored by whether the samples compared were collected from the same hospital (red) or from different hospitals (grey). c) Pairwise ACNI distribution of strains mapping to references representing the five major lineages identified by Shao *et al.* ST179 was not represented by any reference in our database, but the closest match was ST711. d) StrainGE’s ACNI had good correlation with ANI between isolate genomes. e) StrainGE provided strong evidence for maternal strain transmission. f) When including other babies with strains close to *E. faecium* DMEA02 and *E. faecalis* SF28073, same-subject comparisons were firmly in the top right corner with high ACNI and high gap similarity, while the majority of between-subject comparisons were more different.

We also noted that the isolate-based tree topology in Figure 6a varied across the major lineages, with some lineages having more consistent short branching (e.g., ST191) than others (e.g., ST40), indicating clear patterns in relationships among close isolates. The StrainGR-computed ACNI values largely mirrored these results in how narrow (e.g., ST191) or wide (e.g., ST40) the ACNI distributions were for comparisons across predicted members of these lineages (Figure 6c). This correlation was also observed when we examined the relationship between ACNIs, calculated by StrainGR using metagenomic data, and ANIs, calculated by FastANI (25) using isolate genomic data, from the same stool samples (Pearson’s *r*=0.95; Figure 6d; Materials & Methods), though StrainGR consistently reported higher ACNI than ANI.

To determine instances of mother-to-child strain sharing, Shao *et al.* used StrainPhlan (9), which led them to predict 7 *E. faecalis* and 2 *E. faecium* maternal strain transmission events. Though we could not directly compare our results (the sample names for the predicted mother-baby pairs were not reported), we sought to determine whether StrainGE could identify comparable numbers of mother-to-baby strain sharing. StrainGST reported the same reference for 17 mother-baby pairs. Six of these had sufficient common callable genome to assess ACNI between predicted strains; three mother-baby pairs had ACNI near 100% and three had an ACNI <99.7%. In one case, we identified a baby *E. faecium* strain with 99.999% ACNI to a strain in the mother (Figure 6e). Even though many of our strains were closely related (Figure 6b), only 0.9% of comparisons had an ACNI at least that high. When comparing strains from this mother-baby pair to those matching the same reference in other mothers or babies, they generally had considerably lower ACNI and gap similarity (Figure 6f), highlighting StrainGR’s ability to distinguish same-strain relationships. For the other pairs having ACNI <99.7%, StrainGE results from the larger dataset suggest that these strains cannot be confidently discerned from locally circulating strains and, therefore, may not represent mother to baby transmission.

### Discussion

Strain-level variation can determine a bacterial species’ interactions with its host and host microbiota, and can be used as a tool to trace the dynamics and transmission patterns of a species over time and space. The ability to discern strain-level variation from primary specimens—where the species of interest may be at low abundance—can transform our understanding of them, including species population dynamics and ecology. We have shown that our novel tool suite, StrainGE, works directly from primary specimen metagenomes to enable sensitive differentiation and nucleotide-level characterization of individual bacterial strains, at coverages typical of many clinically relevant bacteria (26–28). StrainGE is highly sensitive and uniquely able to disentangle multi-strain mixtures and to determine strain-specific variation. We also showed that this tool provides a substantial advance over previously published tools, which either report only overall consensus SNV profiles for a mixture (8,9), disentangle strain-specific SNVs in a limited set of housekeeping genes (29), or do not offer nucleotide-level resolution (10,11). Importantly, StrainGE was able to shed light on the strain-level dynamics of a patient with Crohn’s disease, including the presence of co-existing strains and strains at timepoints missed by another popular strain-tracking tool. StrainGE similarly revealed the long-time gut carriage of a UTI-causing *E. coli* strain, suggesting that persistence in the gut may be an important mechanism for UTI recurrence. Finally, using metagenomic data from primary stool specimens, StrainGE was able to recapitulate relationships among stool *E. faecalis* observed from isolate whole genome phylogenetic reconstruction, as well as provide strong evidence for transmission of an *E. faecium* strain from mother to child.

StrainGE is designed to be broadly applicable to many bacterial taxa, and includes tools to adapt to any genus or species of interest. For species with many available reference genomes, StrainGST is capable of handling large reference databases. For species that are less well-represented in genome repositories, our benchmarking (Supplementary Results A; Figure 6) showcased that combining StrainGST and StrainGR can return accurate information about strain relationships, even when fewer reference genomes are available. Though we did not benchmark StrainGE on non-bacterial microbes like fungi or on viruses, StrainGE is tunable and has easy to analyze output that should make it straightforward to explore for these applications.

StrainGST’s default database clustering threshold (0.9 Jaccard similarity, corresponding to an approximate ANI of ~99.8%) should work well for most users, but in some cases it may be helpful to optimize the phylogenetic density of a database in order to maximize the extent of strain-level details provided by the tool. For example, comprehensive representation of the phylogenetic diversity of a taxon can be desirable for identifying the nearest reference strains, since these should serve as the best basis for the identification of strain variants and to tease apart mixtures of closely related strains. However, a database containing nearly identical references can pose a challenge for StrainGST, particularly when strains are present at the extremely low coverages at which StrainGST was designed to operate. Sequencing error and metagenomic background sequence can lead to different instances of the same strain matching to different, but closely related, references, complicating comparisons across samples. Therefore, StrainGST includes the ability to cluster input reference genomes based on a user-defined k-mer Jaccard similarity threshold, reducing the redundancy of the database by selecting one reference from each cluster of highly related references. We recommend using assemblies that are as complete and as contiguous as possible in database construction, since these make plasmid detection, read alignment and variant calling more straightforward. However, StrainGE can work with less complete assemblies, though the user should be aware that they contribute to additional sources of error when analyzing StrainGE output.

Using a clustered database, StrainGST’s k-mer-based scoring heuristics (Materials & Methods) are capable of detecting and correctly identifying nearest references of strains with high precision and recall down to 0.1x coverage per strain, or about 0.02% relative abundance, assuming a 5Mb genome with 3 Gb of sequenced reads. StrainGST’s iterative approach to strain identification finds and reports the nearest reference for the strain with the most evidence, and then excludes all k-mers from that reference from consideration in future iterations, thereby focusing on distinct evidence to identify subsequent strains. This has proven very effective in identifying mixtures of strains throughout the phylogeny of both equal and unequal abundance. By using StrainGST alone, the user can get a sense as to which strains of the target taxon are present in a sample, their relative abundances, as well as what known references they are most like, enabling straightforward placement of observed strains into the known phylogenetic structure of the database.

StrainGR provides a deeper level of context beyond StrainGST, identifying genetic differences between the actual strain(s) and the reference(s) selected by StrainGST, by aligning metagenomic reads to the StrainGST-identified references. The combined reference set allows the aligner to place reads at their optimal location, deconvolving reads with alleles specific to a strain. In order to be robust against sequencing and alignment errors, StrainGR uses stringent read filters and ignores regions in the reference with abnormally high coverage. StrainGR requires, by default, at least two high quality reads in order to consider evidence of an allele strong, but it also keeps track of alleles present with less evidence (“weak” alleles). Weak evidence becomes useful if there is other reason to believe a strain variant may be present; *e.g.*, if other samples from the same subject have strong evidence for an allele, it is more likely that weak evidence of the same allele in other samples is real. StrainGR is also unique among similar tools in its ability to identify gaps in coverage, *i.e.,* large deletions in the sample strain relative to the reference. Recombination and horizontal gene transfer is very common in bacterial evolution (15), and patterns of presence and absence of blocks of sequence can provide supportive evidence about the relatedness of strains that is orthogonal to SNVs. Further, missing blocks of sequence can point to potentially important missing gene content relative to the reference genome.

StrainGR stands out in its ultra-sensitivity to detect and compare very low abundance strains in a sample. For example, in a 5 Mb genome, StrainGR can make strong allele calls at ~50,000 sites even when data from a strain covers only 0.1x of a detected reference. Though scant, this is sufficient to estimate the ANI between the sample strain and the reference. However, at 0.1x coverage, the probability that related strains from different samples have sufficient read depth at the same location along the reference to call variants is extremely low. Thus, StrainGR performs well in sample comparison benchmarking when coverages reach at least 0.5x, where the likelihood of capturing variation at comparable reference sites is higher. Though we did not benchmark StrainGE on strains at >10x coverage, we noted that StrainGE worked well on a real metagenomic data set where strains were present at 100x coverage, indicating that the tool should also work well at coverages standardly achieved for isolates.

When comparing strains across multiple samples using StrainGR, we found that it can be useful to further refine the set of concatenated references to be used in alignment. Since StrainGR only calls variants within regions unique to a single reference genome by considering only unambiguously placed reads, the more reference genomes identified by StrainGST across samples, the less unique content there is per reference for StrainGR to call variants against. To address this, we included an option (prepare-ref) to further cluster the set of StrainGST-identified references prior to StrainGR analysis, in order to minimize overlap while providing a consistent set of reference genomes for comparing SNVs and gaps across more strains in a sample set. To guide this additional round of clustering, prepare-ref estimates the amount of shared genome in the combined reference using MUMmer (30). Based on a benchmark with several *E. coli* mixtures (data not shown), we set the default clustering value to a Jaccard similarity of 0.7 (~99.2% ANI), which worked well for our use cases. Application of prepare-ref to the dataset shown in Figure 4 increased the total amount of unique content across references from 19% (including all 14 references) to 57% (including 10 references after collapsing), and enabled direct comparisons of more strains with respect to their nucleotide and gap similarities. This parameter can be easily adjusted by the user.

Despite StrainGE’s focus on unique regions, we found that StrainGE-computed ACNI values were strongly correlated with true ANI. In our *in silico E. coli* benchmark, this correlation was strong even in the context of other diverse *E. coli* and metagenomic backgrounds (Figure 3b). We also found a strong correlation in a real data set comparing ACNI between *Enterococcus* strains to the ANI of isolate genomes from the same metagenomes, even though ACNI was consistently higher than the ANI (Figure 6d). This difference could be because FastANI (25) was used to compare draft isolate assemblies, possibly including comparisons of undetected plasmid contigs, while StrainGE’s ACNI was computed by comparing the chromosome only. This difference could have also arisen due to StrainGE’s stringent read filters, which may have prevented calling true variation in highly divergent regions. Though the measures that define the “same” versus “different” strain will depend upon the research question and the species being evaluated (6), our benchmarking suggested that when both ACNI and the gap similarity between two strains were >99.98% ACNI and >0.97 gap similarity, respectively, this was strong evidence that strains were essentially the same. These thresholds are unlikely to be universal for all studies or species. However, StrainGE provides a compendium of outputs that give the user the ability to assess relationships between strains in detail. While StrainGE’s default settings limit SNV reports to only the unique regions of each reference since it is unable to disentangle variation at multi-mapped sites, there is an option to have it report variation across the entirety of each reference, which could be helpful to characterize variation in known genes.

In summary, StrainGE is an easy-to-use tool for ultra sensitive and high resolution tracking of strains in metagenomes. Rather than relying on a single approach, StrainGE uses both k-mer and alignment analysis to reveal as much information as possible about sample strain genomes, including basic high-level information about strains such as their i) closest matching reference, which places them phylogenetically, ii) relative abundance, and iii) ACNI to other strains, which can be achieved even at very low coverage levels, with more detailed information about specific variants and cross-sample comparisons becoming available as coverage increases. StrainGE is limited in evaluating the relationships between strains using unique regions of reference genomes, is unable to detect new genes that occur in strain genomes that are not present in its closest matching reference, and currently only works with Illumina data. Furthermore, StrainGE is currently not designed to phase SNVs from multiple strains matching the same reference in the same sample. In this case, StrainGR output will reveal evidence for multiple alleles, the frequencies of which cannot be robustly compared to link alleles together at the coverages under which StrainGE was designed to operate. Despite these limitations, StrainGE provides a major advance compared to previously published tools and will help to accelerate our understanding of the role of strain-level variation in shaping ecological and disease processes.

### Materials and Methods

#### Strain Genome Explorer toolkit algorithms

##### StrainGST: Strain Genome Search Tool

StrainGST is a k-mer based tool used to identify specific strain(s) of a species in a metagenomic sample. StrainGST computes a reference database of previously sequenced strains from this species, and uses it to report close reference genomes to strains present in a metagenomic sample along with their relative abundances. The references reported by StrainGST can be used as input to StrainGR to further characterize genetic variation found within the metagenomic sample.

###### Creating a StrainGST database

A StrainGST database is constructed from a set of high quality sequenced reference genomes for a single species or genus, such as all complete reference genomes in NCBI RefSeq. From this set of genomes, StrainGST generates a database of k-mer profiles, using a sliding window (window size k) to traverse each genome and count the frequency of each k-mer. To reduce memory usage and computation time, a minhash technique (similarly to Mash (18)) is applied to keep 1% of the k-mers with the lowest hashes.

StrainGST next performs clustering to remove highly similar genomes from the reference set. In order to track and compare genomic variation across related samples, StrainGR must be able to align reads to a common reference genome across different sample sets. Therefore, the references reported by StrainGST should not be too closely related, or each sample could end up matching distinct yet closely related references, making comparisons difficult. StrainGST computes pairwise Jaccard similarities using each reference genome’s k-mer set, performing single linkage clustering using a Jaccard similarity threshold of τ, and picking a single representative genome for each cluster to include in the reference set. StrainGST selects the genome with the highest mean similarity to all other genomes in that cluster. This process ensures that the k-mer similarities between remaining genomes in the database are all lower than τ. Additionally, to remove genomes from the database that are highly similar to another genome, but that may have lower Jaccard similarity due to the presence of large indels, StrainGST removes genomes where 99% or more k-mers overlapped with those from another genome.

###### Identifying strains present in a sample

StrainGST uses this database to identify the closest reference genome(s) to the strain(s) present within a sample (Figure 1). First, all reads in the sample are k-merized, resulting in the k-mer set *K_sample_*. The algorithm then selects for k-mers from the species of interest by taking the intersection between the sample k-mer set and that of the reference database for the species of interest (Figure 1b), excluding k-mers not associated with the target species.

StrainGST then uses these k-mers to identify the reference genome(s) with the best k-mer matches to the sample using an iterative process. In each iteration, StrainGST scores each reference genome in the database against the remaining k-mers in *K_sample_* in order to find the reference with the best score, which is reported to the user as the reference with the strongest evidence of being present. The scoring system is described in detail below. If no reference strain is identified that scores above a threshold θ(adjustable by a command line option), the algorithm is terminated. The default value for θ (0.02) was optimized to maximize sensitivity while minimizing false positives (Supplementary Results D). In each iteration, k-mers corresponding to the reference selected are removed from the sample k-mer set in order to enable identification of secondary strains in the next iteration. This process continues until either no strain is reported or the maximum number of iterations is reached (default of 5).

###### Scoring metric for selecting matching reference strains

To determine which reference strain to report in each iteration, we calculate a score for each reference strain using a combination of three metrics based on: 1) the fraction of matching k-mers in the reference; 2) the fraction of the reference database k-mers in the sample that could be explained by this reference genome; and 3) the evenness of the distribution of matching k-mers across the genome.

###### 1) Fraction of matching k-mers in the reference (*f*)

This metric represents the fraction of distinct k-mers in reference *j* that is present in the sample and has a value between 0 and 1, where 1 would indicate all k-mers of this reference are present in this sample.

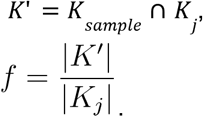

*K_sample_* represents all k-mers in the sample, *K_j_*. represents all k-mers in reference *j*, and *K*’ represents the set of k-mers both present in the reference and in the sample.

###### 2) Fraction of sample k-mers that could be explained by this reference (*α*)

To give more weight to reference genomes that are similar to higher abundance strains in the sample, StrainGST calculates the fraction of database k-mers remaining in the sample that could be explained by the k-mers in this reference:

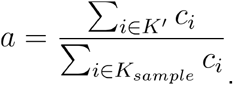

*c_i_* represents the count of k-mer *i* in the sample. Note that we include k-mer counts, rather than using the fraction of *distinct* k-mers, which gives more weight to reference strains with high average depth of coverage. This metric has a value between 0 and 1.

###### 3) Evenness (*e*)

To quantify whether the matching k-mers are evenly distributed across the reference genome, rather than being found predominantly in a small region (*e.g.*, due to a horizontal gene transfer event, or conserved regions attracting reads from different species), we defined the *evenness* score. First, we assumed that the coverage across the genome follows a Poisson distribution. The rate parameter *λ_j_*, of the Poisson distribution specifies the average depth of coverage across the whole genome:

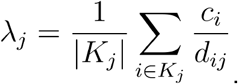

*c_i_* represents the count of k-mer *i* in the sample, and *d_ij_* represents the count of k-mer *i* in reference strain *j*.

If X is the random variable that indicates how many reads cover a position, then using the Poisson distribution, the probability of observing *x* reads at a position is:

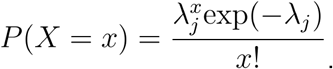

The probability of observing 0 reads at a position is then *P*(*X* = 0) = exp(-*λ_j_*). The probability of observing at least one read at a position is (24):

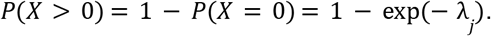

This probability also represents the expected fraction of the genome covered by at least one read given a certain average depth of coverage. The evenness score describes how well the observed fraction of the genome covered by at least one read (which is estimated using the fraction of matching k-mers in the reference defined earlier), matches the expected fraction of the genome covered by at least one read when assuming a Poisson distribution for the depth of coverage:

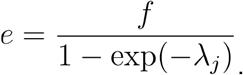

This score will be close to 1 if the observed fraction of the genome with at least one read matches the expected value for a certain average depth of coverage (assuming a Poisson distribution). It will be closer to zero if only small portions of the genome are well covered. A value higher than 1 indicates that the observed fraction of the genome with at least one read is higher than the expected fraction of the genome with at least one read. To bound this score between 0 and 1, StrainGST uses the minimum of *e* and its reciprocal:

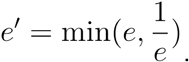

Finally, we combined these three metrics together in order to obtain the final score:

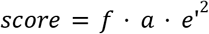

At each iteration, the reference strain with the highest score represents the best match to the highest abundant strain remaining in the sample and is reported to the user.

#### StrainGR: Strain Genome Recovery

The StrainGR pipeline consists of: 1) building a concatenated reference based on reference strains reported by StrainGST; 2) aligning reads to the concatenated reference; 3) analyzing read alignments to called SNVs and large deletions; and 4) using these variant calls to analyze gene content or track strains across multiple samples.

##### Preparing a concatenated reference

To analyze a set of related samples together, such as a longitudinal series, StrainGR concatenates a single, unified set of representative references present across the whole dataset. This can facilitate comparisons of alignments or genomic variation across a set of samples, which may contain different strain mixes at different time points. Use of the concatenated reference allows reads with an allele unique to a particular strain to be aligned to the genome of the correct reference strain, thus helping disentangle reads from mixture samples. Genomes from the same species, however, will share conserved genomic regions (*i.e.*, housekeeping and other core genes), where the aligner will be unable to place reads unambiguously within the concatenated reference. StrainGR detects and excludes these conserved regions from variant calling.

In order to minimize conserved regions where StrainGR is unable to call variants, it is important to select a set of reference strains that are not too closely related, which could result in a large fraction of the concatenated reference genome being marked as shared. To construct a concatenated set of references with an optimal degree of similarity, StrainGR includes a tool called prepare-ref that analyzes StrainGST output from a set of samples (*e.g.*, a longitudinal set from a single patient) and generates a concatenated reference ready for use with StrainGR, optionally performing another round of clustering at a stricter threshold to prevent too-closely related genomes from being included. By default, the stricter clustering threshold is set to a Jaccard similarity of 0.7 (~99.2% estimated ANI).

##### Read alignment and filtering

The reads from a metagenomic sample are then aligned to the concatenated reference using BWA mem (25), removing read pairs with 1) improper pairing; 2) clipped alignment; or 3) implied insert size smaller than the read length. In order to identify shared regions within the concatenated reference which should be excluded from variant calling, StrainGR tracks the number of “multi-mappable” read alignments (those which map equally well at multiple locations) at each position in the reference. When the majority of aligned reads at a position are multi-mappable, StrainGR excludes this position from variant calling. We rely on BWA’s “XA” SAM tag to obtain a read’s alternative alignment locations, so aligners other than BWA are not currently supported by StrainGR.

In addition to excluding multi-mappable regions, StrainGR also excludes regions with abnormally high coverage (greater than threshold τ), likely due to genes highly conserved across genera which attract nonspecific reads from other members of the microbial community. τ was chosen such that the probability of observing a depth of coverage higher than τ is 1×10^-7^ assuming a Poisson distribution. This value results in a threshold of 10x coverage when the mean coverage depth across the genome is 1x, and a threshold of 20x when the mean is 5x.

##### SNV calling

StrainGR analyzes read alignments to identify single-nucleotide variants (SNVs) between a specific strain within a metagenomic sample and its closest reference genome identified by StrainGST. To filter likely sequencing errors, bases with an Illumina Phred base quality score <5 are ignored by default. An allele is considered strong if the sum of base quality scores supporting that allele is i) higher than 50 (roughly equivalent to having at least two high-quality supporting reads) and ii) at least 10% of the total sum of base quality scores of all alleles at that genomic position. If an allele is present but doesn’t match these criteria, it is considered weak. StrainGR stores weak evidence for use when tracking a strain across multiple samples—if a particular strain is highly abundant in some samples, with many strong SNP calls, then weak calls can be useful to discern that allele in low abundance samples from the same sample set.

Based on the observed alleles, StrainGR classifies a genomic position as either “reference confirmed”, “SNV” or “multiple alleles”. If a position has a single strong allele call, and that allele is the same as the reference, the position is classified as “reference confirmed”. A position with a single strong allele call that is different from the reference is classified as a SNV. Any position with multiple strong allele calls (whether they match the reference or not) is classified as “multiple alleles”.

To estimate the overall degree of similarity between the strain in the sample and its closest reference, StrainGR computes an estimate of average nucleotide identity (ANI) using StrainGR SNV calls: the average callable nucleotide identity (ACNI) is the percentage of positions marked as “reference confirmed” out of all positions with a single strong allele call.

##### Large deletion predictions

StrainGR also analyzes the read alignments to identify large deletions present in a specific strain within a sample, as compared to its closest reference identified by StrainGST. Consecutive positions in the reference genome over a specified length (by default 5,000 bp; ~2 genes) with no aligned reads could indicate a large deletion. To account for situations with low coverage across the genome (<1x), StrainGR employs a simple heuristic that exponentially scales the threshold for the length of such regions at lower coverages; thus, only longer gaps can be detected at lower coverages. If λ is the average depth of coverage along the genome, and ϕ is the unadjusted threshold, then the adjusted minimum size of a “gap” is:

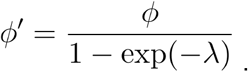

Large deletions are used to assess whether particular genes are absent from the strain in a sample. In addition, the overall pattern of deletions across the genome for the strain in a longitudinal sample set can be used as a strain “fingerprint” to track a particular strain across samples.

##### Strain comparisons across samples

To assess whether the strains in two samples are the same (or very closely related), we compare both SNV calls (via pairwise ACNI) and patterns of large deletions. StrainGR calculates pairwise ACNI by dividing the number of positions where both samples have the same strong allele by the total number of positions where both samples have a single strong allele. To compare the pattern of predicted deletions between two samples, StrainGR calculates the Jaccard similarity: if *G*_1_ is the set of positions *not* marked as a large deletion in sample 1, and *G*_2_ is the set of positions *not* marked as a large deletion in sample 2, then the gap similarity *l* is defined as

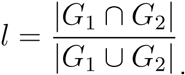

### Benchmarking StrainGE using simulated data and mock communities

#### Spiked metagenome generation

Unless otherwise noted, all synthetic metagenomes used for benchmarking were generated as follows: reads were simulated from the relevant genomes using ART (31) and merged with reads subsampled from a genuine metagenomic data set without detectable *E. coli* (accession SRS014613) as per MetaPhlan2 (19) and StrainGST, up to a fixed depth of 3 Gb. At this depth, strain coverages of 0.1x, 0.5x, 1x and 10x corresponded to relative abundances of 0.016%, 0.083%, 0.16%, and 1.6%, respectively, assuming a 5 Mb *E. coli* genome.

#### *StrainGST database for* Escherichia

For construction of the *Escherichia* reference database, all complete *Escherichia* genomes available in NCBI RefSEq were downloaded in July 2019 (929 genomes total; Supplemental Table 1). All tools required to construct the StrainGST database are included in the StrainGE suite.

In order to set StrainGST’s default clustering threshold, we benchmarked its ability to correctly identify single strain and two-strain mixes using the metagenomic spike-in methods described below, using synthetic reads generated from 200 randomly selected *E. coli* genomes spiked into subsets of real metagenomic samples devoid of *E. coli*, to a total of 3 Gb. For the single-strain benchmarks, 200 samples were generated with 10x, 1x, 0.5x, and 0.1x coverage of each of the selected *E. coli* genomes (800 samples total). For the 2-strain mix benchmarks, 100 random 2-strain combinations from the set of 200 selected *E. coli* genomes were spiked in at each combination of 10x, 1x, 0.5x, and 0.1x coverage (10 coverage combinations, 1000 samples total). The 1800 benchmark cases were run using database clustering thresholds of 0.95, 0.90, 0.85, and 0.80 Jaccard k-mer similarity, corresponding to Mash distance ANIs of 99.89%, 99.77%, 99.63%, and 99.49%, respectively. For each threshold, we measured precision, recall, and F1 score for strain identification, with true positives being only those cases in which StrainGST identified the closest reference strain to the true strain as measured by Jaccard k-mer similarity. The clustering threshold of 0.90 generated the best combined results in each of the three metrics (Supplementary Table 4).

#### Phylogenetics and MLST typing of genomes in the Escherichia reference database

A single copy core (SCC) phylogeny was generated for the entire database of reference genomes. In brief, SynerClust (32) was used to generate clusters of orthologous genes (orthogroups). A concatenated alignment was generated for all single-copy, core orthogroups using MUSCLE (33). A phylogenetic tree was constructed using FastTree v2.1.8 (34). Phylogenetic trees were visualized using iTol (35).

MLST designations for each reference genome were predicted with the tool mlst (https://github.com/tseemann/mlst). Sequence types reported were based on the Achtman scheme. *E. coli* clade/phylogroup designation was determined using ClermonTyping (https://github.com/A-BN/ClermonTyping). For cases when there were missing or conflicting results between predicted typing and MASH groups, the clade designation for a given genome was selected based on where it was located in the SCC phylogeny with respect to unambiguous genome to clade designations.

#### Creation of four-strain E. coli mock community

Four phylogenetically distinct *E. coli* strains - H10407 (clade A), E24337A (clade B1), UTI89 (clade B2), and Sakai (clade E) - were cultured separately overnight at 37°C in 2 mL of liquid LB media shaking at 200 rpm. The bacterial number in each culture was estimated via optical density and then combined at a ratio of 80% H10407, 15% UTI89, 4.9% Sakai, and 0.1% E24337A. Genomic DNA was then extracted from this mock community using the Qiagen MagAttract DNA Isolation Kit (Hilden, Germany), following manufacturer’s protocols. In two separate tubes, human genomic DNA was then added to the extracted *E. coli* DNA for final ratios of 99% human / 1% *E. coli* (weight / weight). Sequencing data for this mock community has been submitted to NCBI’s Sequence Read Archive (SRA) under bioproject PRJNA685748 (biosample SAMN17091845).

#### Comparison of tools for tracking specific strains across samples using simulated sets of related samples

We compared the ability of StrainGE, StrainPhlan (9), and MIDAS (8) to track strains across samples. We performed strain tracking comparisons across ten sets of twelve spiked metagenomes, where each set of twelve was structured similarly in terms of strain content (Figure 2a-b). For each set, we randomly picked two *Escherichia* reference genomes (A and B) from NCBI RefSeq complete, and derived two different but closely related synthetic strains from each reference by introducing ~5,000 random SNVs (99.9% ANI) uniformly across the genome. We spiked reads generated from these synthetic genomes into a real metagenome to generate samples containing these strains in different combinations (Figure 2b), at 0.1x, 0.5x, 1x and 10x coverage.

For each data set, we assessed strain similarity metrics calculated by each tool, to determine whether the tool could match i) the identical strain found in different samples (*i.e.,* strain A in sample 1 and 2; Figure 2c); ii), strains found either in mixtures or single-isolate samples (*i.e.*, strain A in sample 1, 2, 5, and 6; Figure 2d); or iii) closely related strains (*i.e.*, the ability to distinguish strain A1 from strain A2; Figure 2e). In each case, we compared the tools’ predictions to the known strain content of each sample to calculate true positives (TP), false positives (FP), and false negatives (FN). For each tool, we varied the threshold (discussed in detail below) for determining shared strains in order to plot precision-recall curves.

##### Detecting shared strains using StrainGE

For each sample, we ran the complete StrainGE pipeline: StrainGST was run to identify the closest reference genomes, and StrainGR was run on a sample-specific concatenated reference to call genetic variation. To detect shared strains, we collected all samples predicted to match to the same StrainGST reference, and computed a pairwise ACNI matrix for strain comparisons with at least a 0.5% callable genome. The similarity matrix was transformed to a distance matrix by computing 1 – *ACNI*, and transformed to a genetic distance using the Jukes-Cantor model (36). If a pair of samples did not share any predicted close reference genomes, we set the distance between those samples to the maximum integer value. To plot the precision-recall curve, we varied the genetic distance threshold that determines when strains are considered the same.

##### Detecting shared strains with StrainPhlan

We ran StrainPhlan on each sample, using the tool’s marker gene database v295 (Jan 2019). Using the marker gene SNV profiles for each sample, StrainPhlan computed the pairwise sample distance matrix using Kimura’s two parameter model (37) (as suggested in their user manual). To plot the precision-recall curve, we varied the genetic distance threshold that determines when samples share a strain, as performed for StrainGE. To tune StrainPhlan for lower coverage levels, we ran it using `--relaxed-parameters`.

##### Detecting shared strains with MIDAS

We ran MIDAS v1.3.2 (database version v1.2) with default parameters. MIDAS includes a strain tracking tool that is first “trained” by giving it a single sample from each patient in a cohort. This training step identifies unique SNV markers for each patient. For our benchmarking, we “trained” MIDAS on samples containing a single strain (sample 1 for strain A1, sample 3 for strain B1, sample 7 for strain A2, and sample 9 for strain A2). (This likely helped the tool in benchmarking since, in a real world scenario, it is likely unknown whether a training sample contains a single strain.) Next, MIDAS compares these SNV markers to alleles in other samples and assesses how much they overlap. To plot precision-recall curves, we varied the percentage of overlapping markers between two samples that serves as a threshold to determine whether two samples share a strain. To tune MIDAS for lower coverage levels we ran its `merge_snvs. py` script with `--all_snvs --all_samples` and its `strain_tracking. py` script with `--min_reads 1`.

#### Evaluating the ability of StrainGR to quantify strain sharing in distinct metagenomic backgrounds

In order to determine how well StrainGE metrics recapitulated genetic relationships between strains, we generated another set of spiked metagenomic samples, spiked with varying quantities of *E. coli* reads from real, previously sequenced isolates. Ten stool metagenomes were randomly selected from the Human Microbiome Project (4) (Supplemental Table 3). The randomly selected samples contained *E. coli* at relative abundances between 0.005% and 0.9%; no two samples contained the same *E. coli* strain based on StrainGST output. Ten complete genome sequences of *E. coli* isolates, distinct from those identified in the background metagenomes, were selected from NCBI RefSeq database. For each isolate, ten variants were created by generating random mutations, such that the ANI to the original reference ranged from 99.9% to 99.99% at increments of 0.01%. Each reference and variant (110 in total) were spiked into at least two randomly chosen distinct metagenomic backgrounds at coverage levels of 0.1x, 0.5x, 1x, 2x or 5x. A total of 300 synthetic samples were generated, with 350 pairs containing an identical strain in a distinct background. All spiked samples were analyzed with StrainGE; all sample pairs with a matching StrainGST reference were compared using StrainGR. StrainGST hits corresponding to strains present in background samples were not considered further. The ACNI was calculated for every pair.

### Evaluation of StrainGE on longitudinal, clinical metagenomic samples

#### Metagenomic time series from a patient with Crohn’s disease

We downloaded from the UCSD Qiita database (https://qiita.ucsd.edu/; Supplementary Table 2) 27 metagenomic data sets representing stool longitudinally collected from a single individual with Crohn’s Disease (22). We ran the full StrainGE pipeline on each sample, using our *Escherichia* database and default parameters, to identify and analyze *E. coli* strains. For pairwise strain comparisons, we only included samples where the common callable percentage of the genome was >0.5%.

#### Metagenomic sequencing of longitudinally collected stool

Twelve longitudinally collected stool samples were extracted with Chemagen Kit CMG-1091 (Baesweiler, Germany). Libraries were generated with NexteraXT (Illumina, San Diego, CA, USA) and sequenced in paired-end mode on an Illumina HiSeq 2500 (101 bp length) and/or Illumina HiSeq X10 (151 bp length). Short-read sequencing data was submitted to the Sequence Read Archive (SRA) with Bioproject accession PRJNA400628. We ran the full StrainGE pipeline on each sample, using our *Escherichia* database and default parameters, to identify and analyze *E. coli* strains. For pairwise strain comparisons, we only included samples where the common callable percentage of the genome was >0.5%.

#### Characterization of Enterococcus strain diversity across a large cohort of babies

We downloaded 1679 metagenomes published by Shao *et al.* (24) from ENA (accession number ERP115334). We ran StrainGE on each sample, using our *Enterococcus* database. To compare StrainGE’s ACNI to true ANI between isolate genomes, we downloaded all isolate samples tagged with either *Enterococcus faecalis* or *Enterococcus faecium* from ENA (accession number ERP024601). We *de novo* assembled isolate genomes by first trimming reads with Trimmomatic (38) and then running Spades (39). Assembly QC was performed using GAEMR (http://software.broadinstitute.org/software/gaemr/). Contigs classified as a genus other than *Enterococcus* were removed. To make the ANI computation more comparable to StrainGE’s ACNI, which is based on the chromosome only, we used MOB-suite (40) to detect and remove known plasmids. This removed on average 71 kbp (+/- 64kbp) per assembled genome. We used FastANI (25) to calculate ANI between isolate genomes. Only pairs where FastANI was able to map at least 25% of fragments were considered. For pairwise strain comparisons using StrainGR, we only included pairs with a common callable genome higher than 0.5%. To be able to link ANI between isolate genomes and the correct strain ACNI in the metagenome we ran StrainGST on isolate samples too, to know which chromosome in the concatenated reference to inspect.

## Declarations

### Availability of data and materials

The data generated and analyzed for this publication is available in SRA, under BioProject accession PRJNA400628 (patient with recurrent UTI), and PRJNA685748 (four *E. coli* strains mock community).

StrainGE is an open source tool written in Python and C++ released under the BSD 3-clause license, available at https://github.com/broadinstitute/strainge (10.5281/zenodo.4697575). StrainGE is built on top of several existing Python libraries, including NumPy (41) and SciPy (42). Analysis scripts to generate the figures in this publication are available at https://github.com/broadinstitute/strainge-paper (10.5281/zenodo.4538994).

### Ethics approval and consent to participate

An adult volunteer with a history of UTI was recruited at Washington University, St. Louis under an IRB-approved protocol (principal investigator, Hing Hung (Henry) Lai). Informed consent was obtained.

### Consent for publication

Not applicable

### Author contributions

BJW, CAD, TA, and AME conceived the tool. LvD, BJW, and TJS built the StrainGE software. LvD, BJW, TJS, CJW, AG and CA performed benchmarking analyses. LvD and TJS analyzed real-world metagenomic datasets. HLS, AJP and SJH created and provided essential reagents. LvD, BJW, TJS, AG, ALM, TA, and AME drafted the manuscript. All authors edited and approved the manuscript.

## Acknowledgments

We would like to thank Christopher Desjardins, Ted Pak, Wen-Chi Chou, Rauf Salamzade and Ryan Bronson for helpful discussions.

