## Supplemental Figures and Text for "StrainGE: A toolkit to track and characterize low-abundance strains in complex microbial communities"

### Supplementary Figures

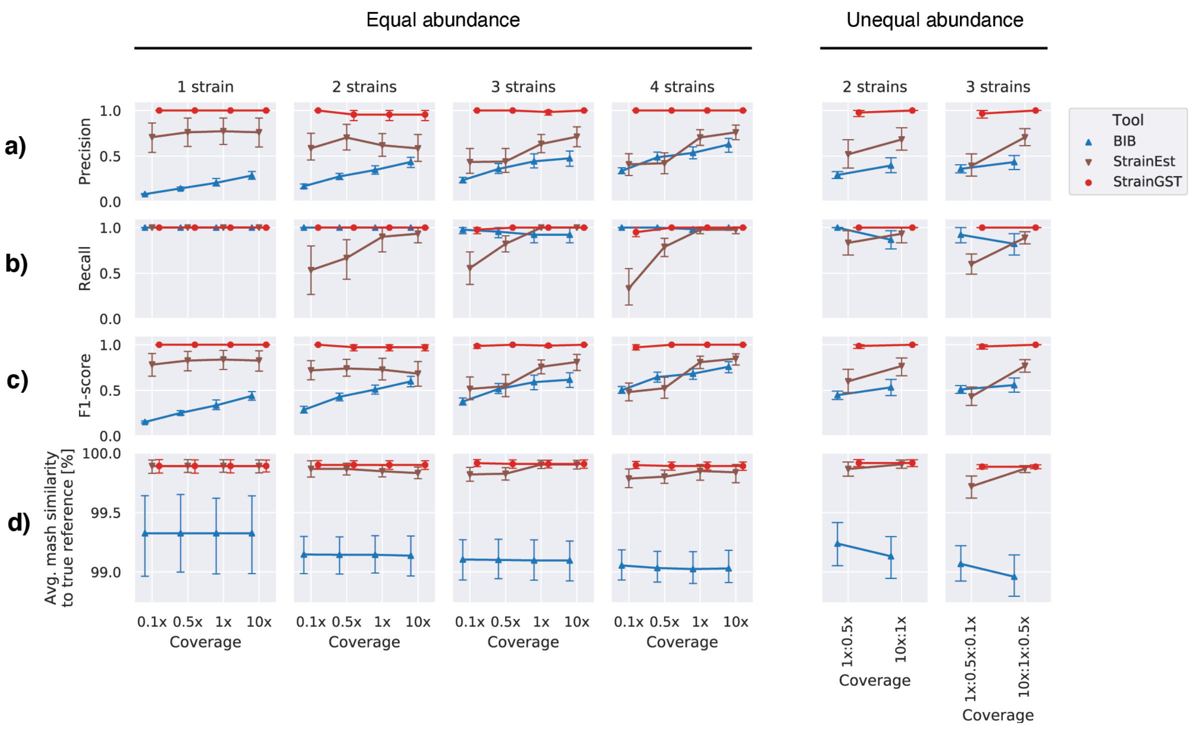


**Supplementary Figure 1. StrainGST was more sensitive and precise in identifying close reference genomes than other tools.** Performance of StrainGST (red circles), StrainEst (brown triangles) and BIB (blue triangles) on 15 sets of metagenomes spiked with known *Escherichia* strains mixed at either equal abundance (left panel; 1-4 strains for each sample, 0.1x-10x coverage) or unequal abundance (right panel; 2 strains mixed at 1x:0.5x and at 10x:1x, or 3 strains mixed 1x:0.5x:0.1x and 10x:1x:0.5x). Performance plotted as a) Precision, b) Recall, c) F1 score, d) Average Mash similarity to closest reference.

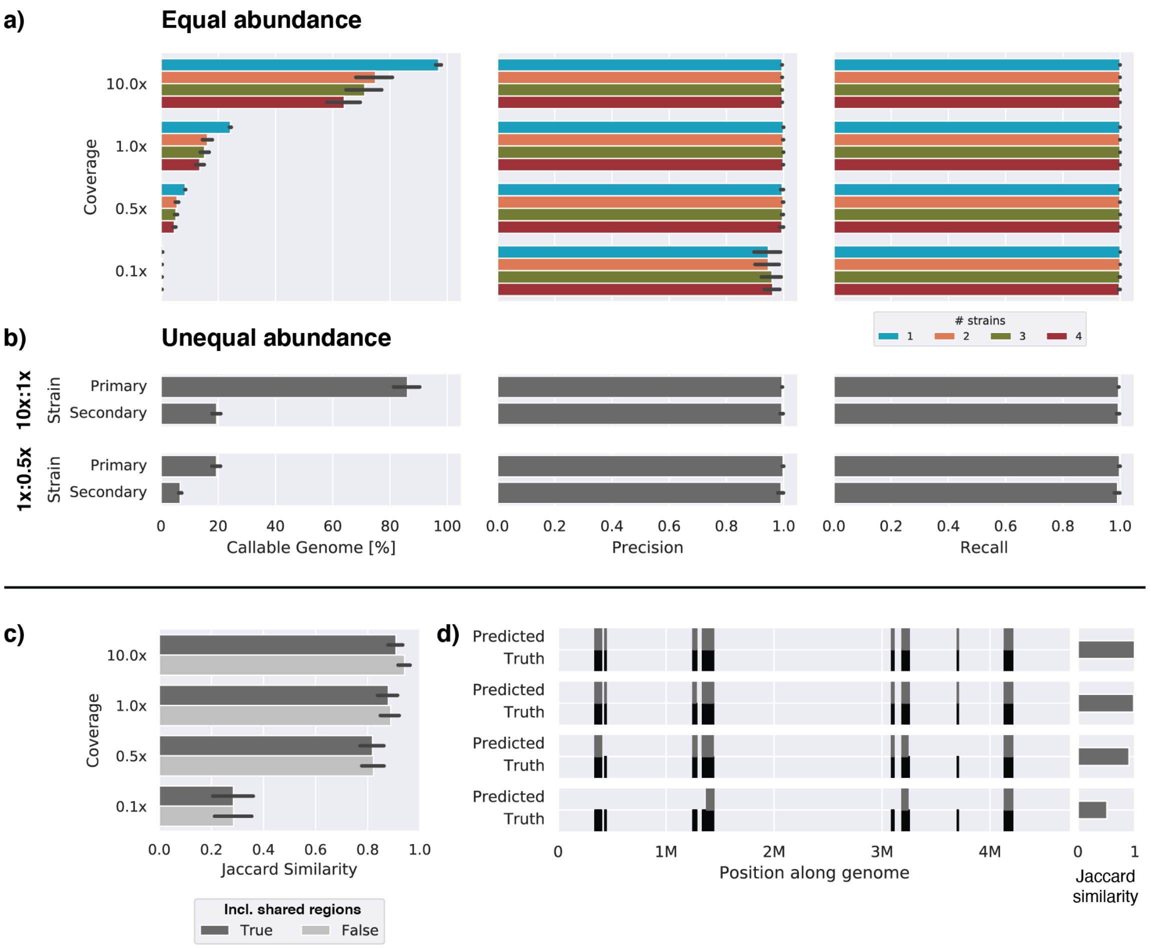


**Supplementary Figure 2. StrainGR accurately called SNVs and large deletions in both single strain and mixed samples at low coverages. a)** Percent callable genome, precision, and recall for SNVs called by StrainGR on mixtures of 1-4 synthetic genomes spiked into a metagenomic sample at different coverages. The “% callable genome” refers to the fraction of the genome where StrainGR was able to make calls. b) Percent callable genome, precision, and recall for SNVs called by StrainGR (limited to callable genome) on pairs of *Escherichia* strains mixed at unequal abundance (1x:0.5x or 10x:1x). c) Jaccard similarity between gaps predicted by StrainGR and known gaps, at different coverages. Dark grey bars indicate the Jaccard similarity when using the whole genome; light grey indicates the Jaccard similarity when ignoring positions with a majority of multi-mapped reads. d) An example of the pattern of deletions present within a synthetic genome (“truth”; black), compared to the pattern of deletions predicted by StrainGR (“Predictions”; grey).

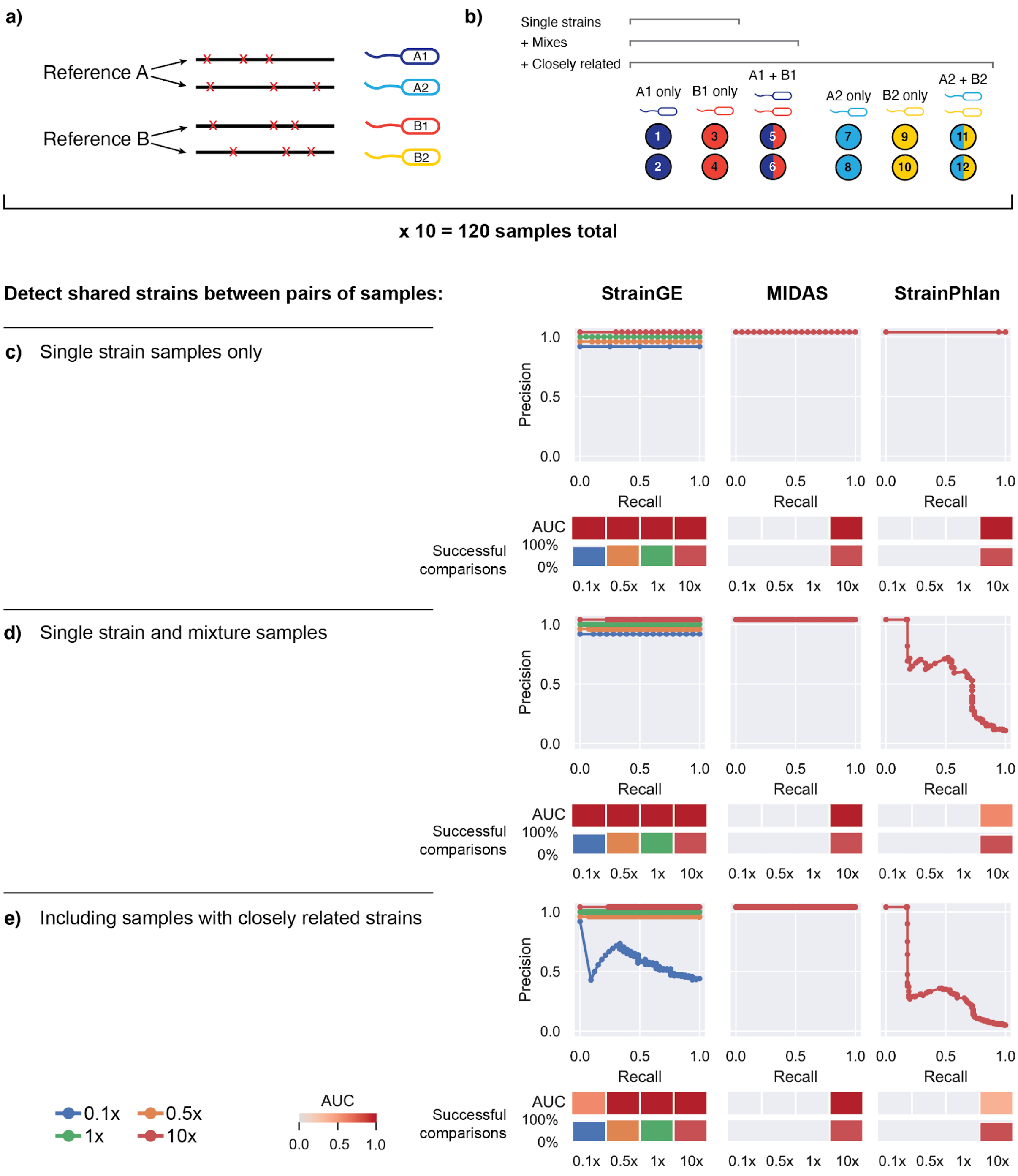


**Supplementary Figure 3. StrainPhlan and MIDAS did not run to completion at coverages lower than 10x with their default settings.** a) Depiction of how synthetic *Escherichia* genomes were generated from randomly selected NCBI RefSeq genomes to create sets of closely related strains (e.g., A1/A2 and B1/B2) for spike in experiments. b) Depiction of how spiked metagenomes were created using synthetic genomes from (a). Each circle represents a spiked metagenome. The color of the circle indicates which synthetic strain was included: single color circles indicate spiked metagenomes containing a single synthetic strain, and two color circles indicate spiked metagenomes containing two synthetic strains mixed at equal proportions. c-e) Precision-recall curves for each tool and coverage 0.1x-10x, when given the task to detect which sample pairs contain identical strains. The area under the curve (AUC) is depicted as a heatmap below. The “successful comparisons” bar plot indicates the percentage of sample pairs for which a comparison was possible (*i.e.*, tools ran to completion for both samples). c) Limiting to single-strain samples from distinct references. d) Including samples with two strains, but limited to strains from distinct references. e) Including samples with closely related strains.

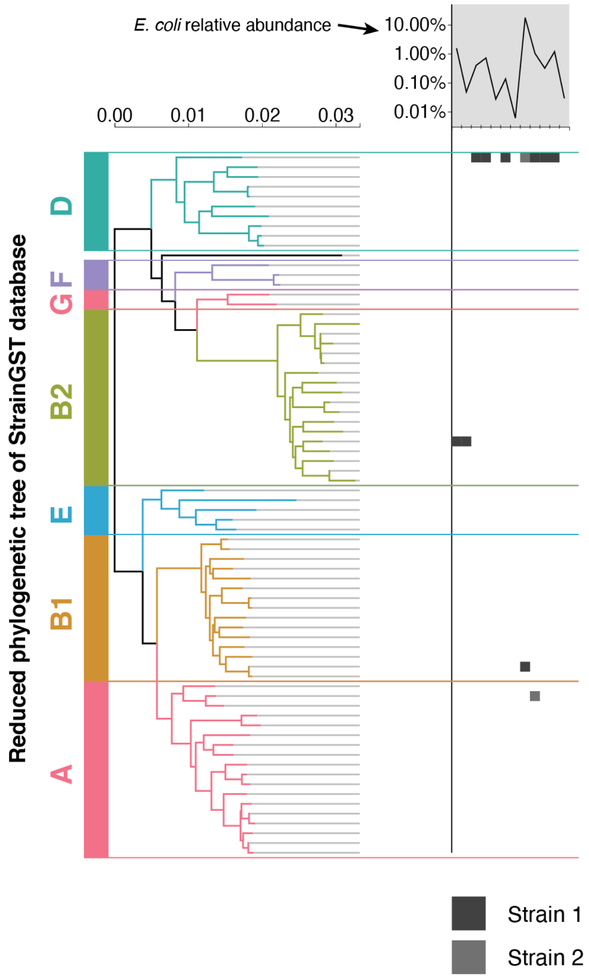


**Supplementary Figure 4. Strains detected by StrainGE can easily be placed in the phylogenetic context of the StrainGST database.** Left panel: reduced phylogenetic tree of the *Escherichia* StrainGST database, with clade designations as obtained using ClermonTyping. Right panel: strains identified from longitudinally collected stool samples from a single individual. Each column represents a time point, and a square within a column indicates the StrainGST match(es) in that sample. Top: overall *E. coli* relative abundance over time.

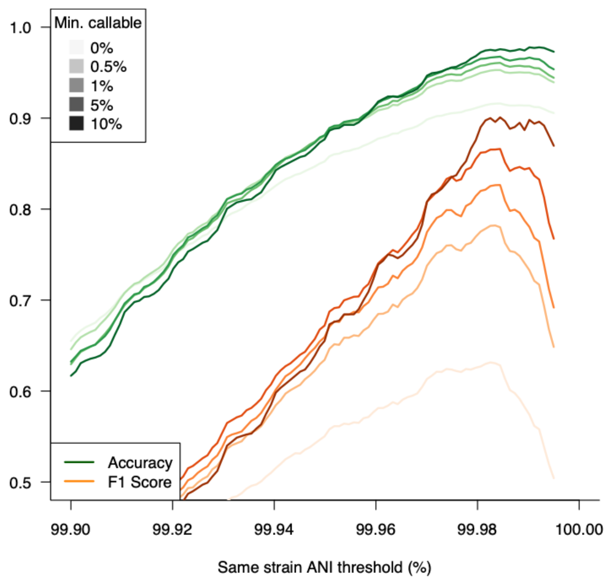


**Supplementary Figure 5**. **StrainGR metrics can be used to accurately classify strain sharing in distinct metagenomic backgrounds.** Using metagenomic samples spiked with *E. coli* isolates containing synthetically introduced SNVs, we used different values for StrainGR's ACNI metric to classify sample pairs as containing the same, or different, strains. All pairs with ACNI above the threshold were considered 'shared' between samples. Pairs were considered correctly classified if the true ANI was 100%. Accuracy (green) and F1 score (orange) were calculated for a range of ACNI thresholds, additionally filtering for comparisons with a minimum amount of common callable genome (light to darker lines). StrainGR's ability to correctly delineate identical strain pairs increased with a larger common callable genome, with a substantial drop in accuracy with common callable genome <0.5%.

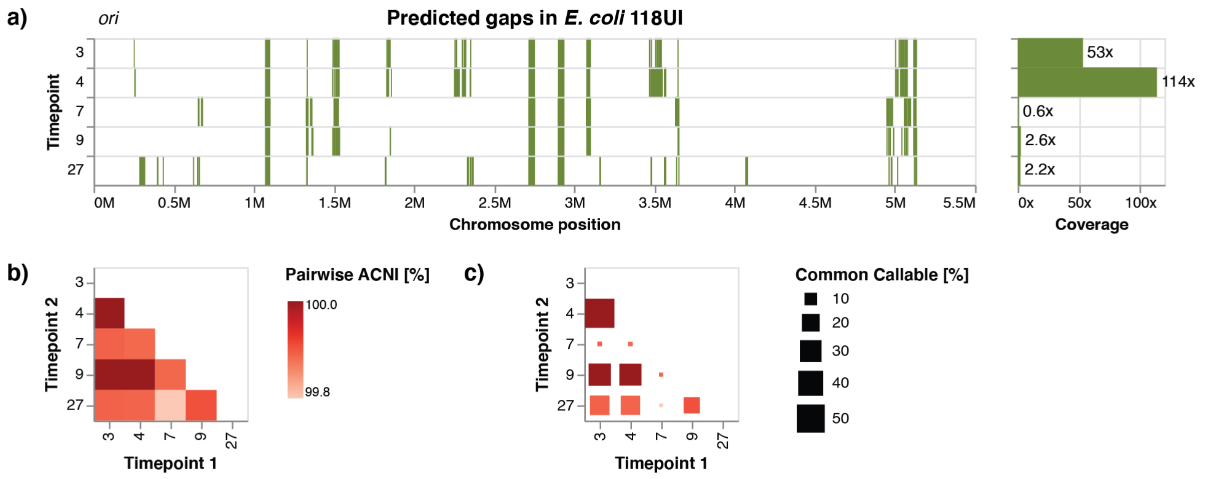


**Supplementary Figure 6. StrainGR provides detailed insights in the genomic diversity within a set of strains close to *E. coli* 118UI.** a) Predicted large deletions in the reference *E. coli* 118UI at multiple time points. Each green rectangle represents a large deletion. b) Pairwise ACNI between strains at different timepoints. c) The same heatmap as in (b), but each square is scaled by the percentage of the common callable genome.

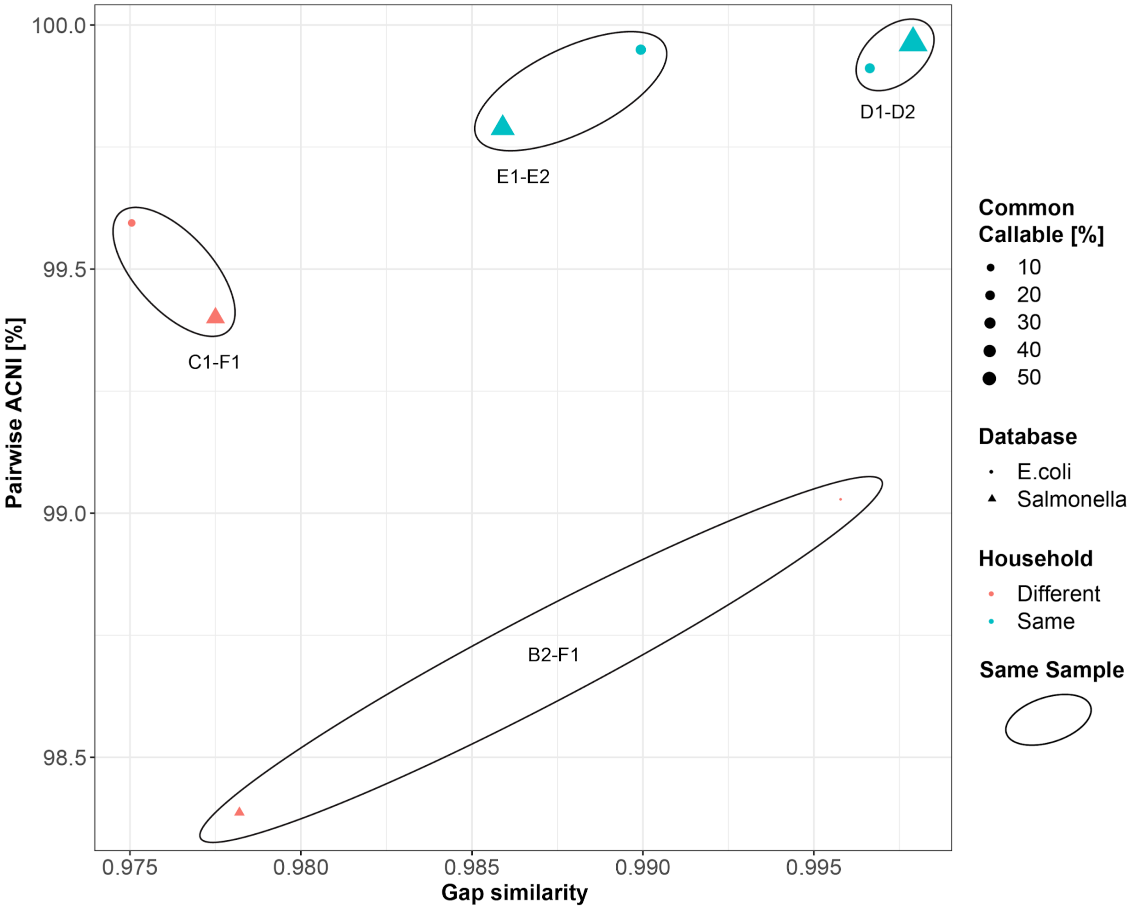


**Supplementary Figure 7. StrainGE can robustly report on strain relationships even when the best match is from a sparse region of the database.** Pairwise ACNI and gap similarity as reported by StrainGE are plotted for the Kenyan household samples which share *E. coli* strains using the *E. coli* database (circle) or the contaminated *Salmonella* database (square). Using the “sparse”, or contaminated, *Salmonella* database, StrainGE is still able to discern close strain relationships (same-household comparisons, teal) from more distant ones (different households, orange), almost as well as when the *E. coli* database is used. Point size reflects the common callable genome between the samples.

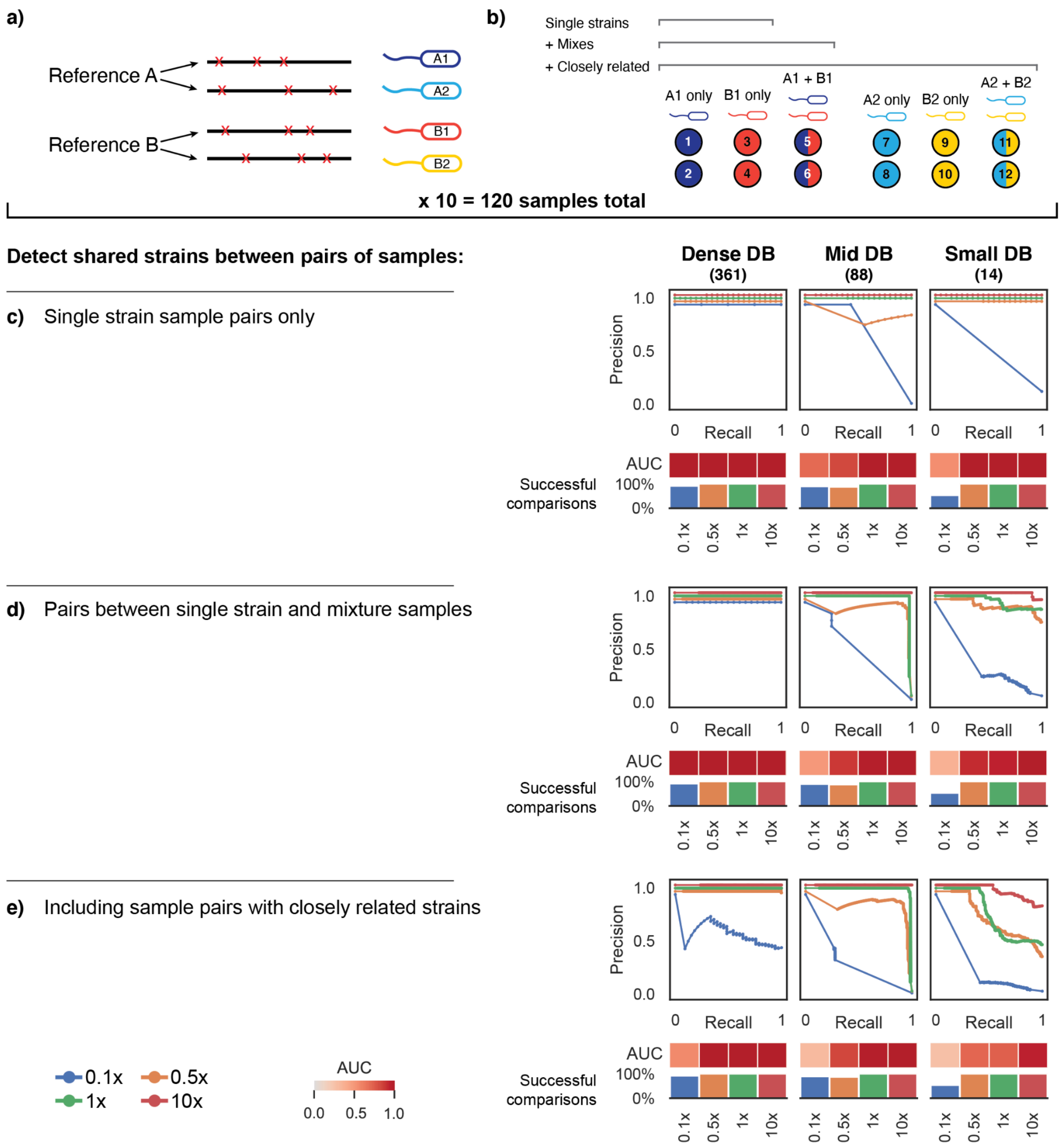


**Supplementary Figure 8. StrainGE could accurately track strains across samples with smaller databases.** a) Depiction of how synthetic *Escherichia* genomes were generated from randomly selected NCBI RefSeq genomes to create sets of closely related strains (e.g., A1/A2 and B1/B2) for spike in experiments. b) Depiction of how spiked metagenomes were created using synthetic genomes from (a). Each circle represents a spiked metagenome. The color of the circle indicates which synthetic strain was included: single color circles indicate spiked metagenomes containing a single synthetic strain, and two color circles indicate spiked metagenomes containing two synthetic strains mixed at equal proportions. c-e) Precision-recall curves for each tool and coverage 0.1x-10x, when given the task to detect which sample pairs contain identical strains. The area under the curve (AUC) is depicted as a heatmap below. The "successful comparisons" bar plot indicates the percentage of sample pairs for which a comparison was possible (*i.e.*, tools ran to completion for both samples). c) Limiting to single-strain samples from distinct references. d) Including samples with two strains, but limited to strains from distinct references. e) Including samples with closely related strains.

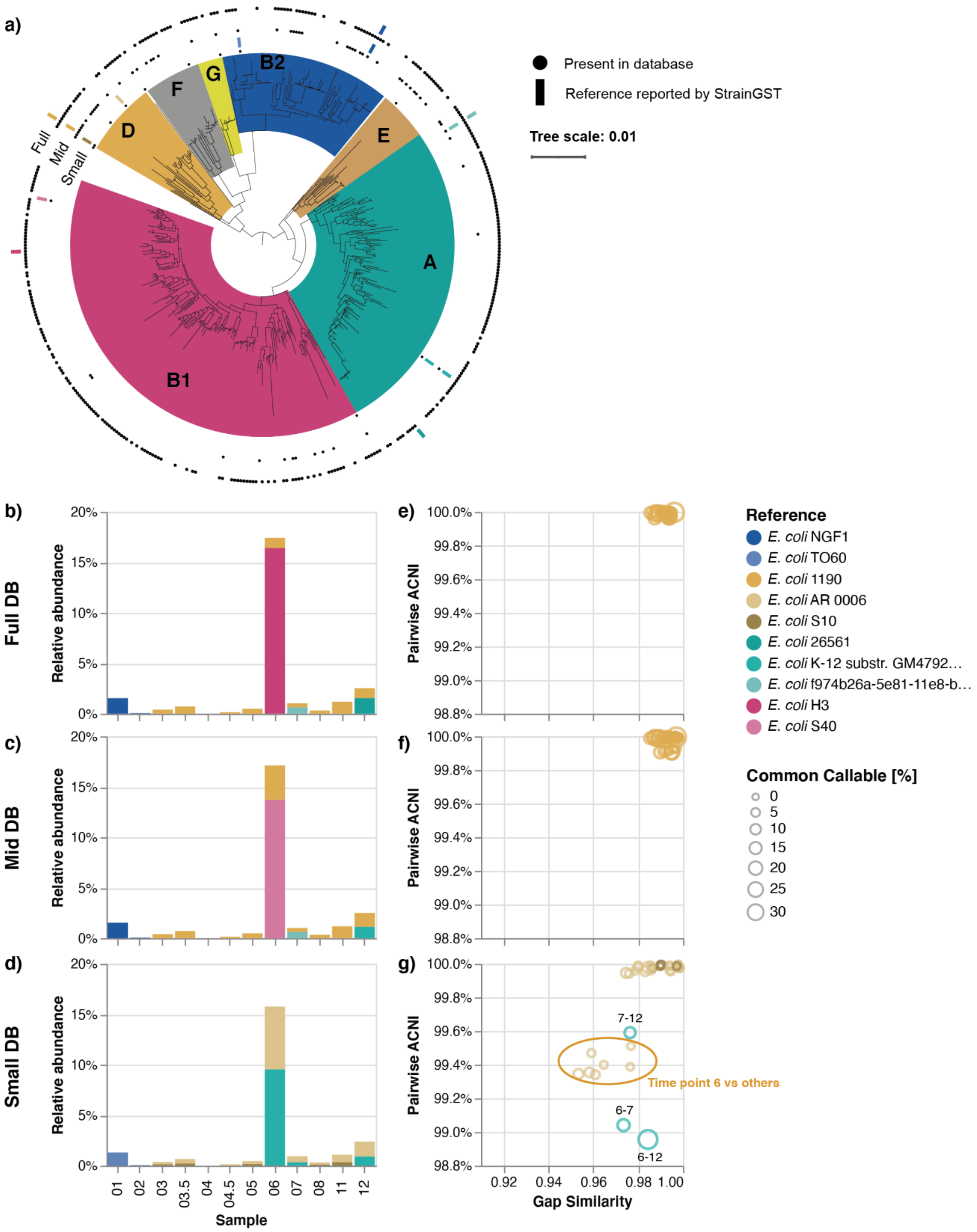


**Supplementary Figure 9. StrainGE provides useful information using both small and large databases on metagenomic samples from a woman with rUTI.** a) Single-copy core phylogenetic tree of 471 *E. coli* genomes (including the full set of genomes contained in the large reference database) with major phylogroups annotated. Black dots indicate presence of the corresponding genome in the small (inner ring), mid and full database (outer ring). Colored rectangles indicate the genomes reported by StrainGST when using the small, mid or full database. StrainGST reported references and their estimated relative abundances per time point when using the b) full database, c) mid database d) small database. For strains matching the same reference, pairwise ACNI and gap similarities are plotted when using the e) small database, f) mid database and g) full database.

### Supplementary Text

#### Supplementary Results

**A. StrainGE can robustly report on strain relationships with sparse databases**.

I. We observed that StrainGE was able to produce reliable results about strain relationships, even when the detected reference strain turned out to be a more distant contaminant in our sequence repository; *i.e.*, from a genetically sparse region of the database. Using a *Salmonella* database, we applied StrainGE to 14 stool microbiome samples obtained from multiple pairs of cohabiting Kenyan siblings, predicted by Kraken2 (1) to contain low levels of *Salmonella* (Supplementary Methods). StrainGST reported that 12 of the 14 samples harbored a match to the same *Salmonella* strain, *Salmonella sp.* HNK130, suggesting that all sibling and non-sibling pairs were colonized with highly similar *Salmonella,* despite households being geographically separated.

Closer examination of StrainGR output revealed, however, that each predicted HNK130-like strain had a different ACNI score relative to the reference, ranging from 98.2% to 99.7%, which were all lower than expected. Subsequent ANI and BLAST analyses (Supplementary Methods) revealed that HNK130 was actually *E. coli*. Recent work has shown (2) that reference genome databases used by Kraken contain cross-species plasmids, like those shared by *E. coli* and *Salmonella*, which can lead the tool to incorrectly assign species. To avoid this problem, StrainGST does not include plasmid content in its reference database, and the tool now flags the user when a reference shares less than 90% ANI to other references.

After removing outliers from the *Salmonella* database, we reran the Kenyan samples through StrainGE, which predicted that there were no *Salmonella* in these microbiomes. Remarkably, the output from running StrainGE on the Kenyan samples using our *E. coli* database provided very similar views of the relative closeness of *E. coli* strains in cohabiting siblings as observed in our original run using the *E. coli* contaminated *Salmonella* database (Supplementary Figure 7). This suggested that StrainGE can return accurate and consistent information about strain relationships, even when the best matching strain is from a low density area of the reference database, which could occur for species of interest that are under-represented in genome repositories.

II. While the above results suggested that our current default threshold for building the StrainGST database could be looser to make the database more sparse, we predicted that there would be trade-offs including potentially losing the ability to resolve more closely related strains and reducing the amount of genome that could be analyzed, given the known relationship between ANI and gene content similarity for many species i.e., more distantly related isolates tend to share fewer genes (3).
In order to formally test this, we first created *Escherichia* databases with increasingly fewer references by clustering the downloaded *Escherichia* references at Jaccard similarity thresholds of 0.8, 0.7, 0.6 and 0.5 (estimated ANI 98.2%-99.5%), resulting in databases with 213, 88, 42, and 14 *Escherichia* references, respectively. Then, to assess the impact of having fewer references on StrainGST's accuracy, we benchmarked these sparser databases against the original denser database (361 genomes; Jaccard

similarity threshold of 0.9) using the same set of 1,800 samples used to optimize the clustering threshold for *E. coli*, each containing 1-2 strains (Materials & Methods). Recall remained consistently high, indicating that StrainGST was able to pick the closest references in the sparser databases. The overall F1-score, however, decreased for smaller databases, driven by an increased number of false positives (Supplementary Table 5). As the closest reference could be quite distant from the sample strain, it was often unable to explain significant portions of the sample genome, leaving sufficient k-mers remaining for StrainGST to report an additional false positive reference.

**Supplementary Table 5. StrainGST performance using various sized databases.**

| Threshold (#refs) | True Positives (TP) | False negatives (FN) | False Positives (FP) | Recall | Precision | F1 |
|---|---|---|---|---|---|---|
| 0.9 (361) | 2721 | 57 | 57 | 0.979 | 0.979 | 0.979 |
| 0.8 (213) | 2640 | 119 | 122 | 0.957 | 0.956 | 0.956 |
| 0.7 (88) | 2532 | 159 | 289 | 0.941 | 0.898 | 0.919 |
| 0.6 (42) | 2284 | 336 | 694 | 0.872 | 0.767 | 0.816 |
| 0.5 (14) | 2138 | 120 | 757 | 0.947 | 0.739 | 0.830 |

To investigate how database sparseness impacted tracking strains across samples, we repeated the *in silico* strain tracking tests (Figure 2) using StrainGR with two smaller databases, clustered at thresholds of 0.5 (very sparse; 14 *Escherichia* genomes) and 0.7 (intermediate; 88 *Escherichia* genomes). As compared to the original database of 361 references, the intermediate database performed comparably for all tests at coverages of 1x and higher (Supplementary Figure 8c,e,d); however, at low coverages (0.5x), a single sample set where StrainGST made incorrect reference calls led to a lower area under the precision-recall curve (AUC) across all tests. For the very sparse database, StrainGE was able to correctly detect shared strains across single strain samples (Supplementary Figure 8c), as well as mixes (Supplementary Figure 8d), at strain coverages of at least 0.5x; however, at low coverages (0.1x) its performance dropped considerably.  In some low-coverage cases, StrainGE was not able to run to completion due to the default minimum coverage requirements not being met (as indicated by the lower "successful comparisons"; Supplementary Figure 8c,d,e). In cases where it did complete, the scant read data aligned less accurately to the more distant reference, which could share as little as ~98% ANI with the sample strain. In addition, this very sparse database also performed worse than denser databases in distinguishing between closely related strains across all coverages (Supplementary Figure 8e), likely because less accurate alignments resulted in lower and less accurate ACNI values.

To examine how the sparser databases would affect results from running StrainGE on real data, we reran the pipeline on the same metagenomic dataset as in Figure 5 (woman with recurrent urinary tract infection) using the intermediate and very sparse databases and compared results to those using the original database. StrainGST results were similar across all databases, with nearly identical read-outs of overall *Escherichia* relative abundance, and reported references mostly from the same phylogroups at similar relative abundances (Supplementary Figure 9b,c,d; phylogenetic distribution of reported references shown in Supplementary Figure 9a). We only observed discordant results with the very sparse database: a strain originally represented by a reference from phylogroup B1 was now represented by a reference from phylogroup A (time point 6); and a phylogroup D strain close to *E. coli* 1190 in the dense database was represented by two phylogroup D references (time points 3-5, 8, 11).

Despite some StrainGST-level differences across database runs, StrainGR was still able to distinguish between "same" and "different" strains that hit the same reference. For example, StrainGR pairwise comparisons of a phylogroup D strain identified by all three (dense, intermediate and very sparse) databases consistently revealed high ACNI and gap similarity, indicating that the strain was the same across samples (Supplementary Figure 9e,f,g). In contrast, for a pair of strains originally mapping to different references with the dense and intermediate databases, but mapping to the same reference with the very sparse database, StrainGR correctly identified that these two strains were not the same, as suggested by the lowered ACNI (Supplementary Figure 9g; time points 6-7).

Though "same" versus "different" strain assignments were generally consistent across databases, we observed a notable difference in ACNI estimates using the very sparse database for comparisons involving time point 6, a sample with relatively high abundance of *E. coli* and consistently predicted to carry two strains (Supplementary Figure 9g). While we can not confirm why ACNI estimates differed so dramatically for the very sparse database, we hypothesize that the reported references in the medium and full database were better representations of the strains in the sample, able to attract reads to the correct locations which improved deconvolution of a strain mixture, resulting in more accurate ACNI values.

In conclusion, while StrainGE can provide useful information even with a small database (still able to pick the closest references), the accuracy of ACNI improved as the database size got larger. Thus, when using a very sparse database and in case of a strain mixture, we encourage users to be more careful interpreting ACNI,, and use any available longitudinal information to confirm the presence of a "same" or "different" strain.

**B. StrainGST works at lower coverages and pinpoints more closely related references than other tools.**

In order to assess the sensitivity and specificity of StrainGST compared to similar tools, we constructed *in silico* metagenomes that were spiked with sequences of known strains of *Escherichia* at varying relative abundances. We simulated reads from randomly selected *Escherichia* genomes downloaded from RefSeq, approximately one third of which were also represented in our *Escherichia* reference database, and mixed them with reads from a metagenomic sample from the Human Microbiome Project without any detectable *Escherichia*, to a total of 3 Gb per sample (Materials & Methods). Strains were mixed at both equal (1-4 strains) and unequal (2-3 strains) abundances to achieve between 0.1x and 10x depth of *Escherichia* coverage (roughly 0.02% to 1.6% relative abundance) per sample per strain, designed to cover the typical ranges of complexity and abundance of *E. coli* within metagenomic stool samples. A total of 240 spiked metagenomes were generated with strains mixed at equal abundance, and another 30 with strains mixed at unequal abundance.

We compared StrainGST to two similar tools, which also identify strains in a sample based on those in a reference database: BIB (4) and StrainEst (5). BIB applies a Bayesian model to sample reads aligned to a core alignment of its database in order to identify the closest strain(s). StrainEst applies a regression model based on unique SNVs in the genomes of strains represented in its database to identify the closest strain(s) in a sample. All tools were run on each spiked metagenome sample, and the fidelity of the results were determined by comparing the reports from each tool against the known composition of the spiked metagenomes. We excluded PathoScope (6) and Sigma (7), because we were unable to run them to completion because of their dependencies on outdated databases or software (Materials & Methods). Because BIB's database construction step, which required generation of a core alignment using progressiveMauve (8), could not scale to include all 361 reference genomes used for benchmarking StrainGST and StrainEst, we also created a smaller database for BIB with only 20 reference genomes.

StrainGST performed as well as, or better than, the other tools across all scenarios tested, and stood out strongly when strains were at very low abundance, either alone or as part of a mixture with other strains. StrainGST had the highest precision (mean 0.99), F1 score (mean 0.99), and its recall and average Mash similarity (9) were at least as good as that of the other tools when given the task of identifying the closest reference(s) to those present in the spiked metagenome (Supplementary Figure 1). There was no significant correlation between StrainGST's F1 scores and the number of strains with an exact reference match in the database (Spearman's $\varrho$=0.06, p-value=0.25) suggesting that StrainGST performance was not dependent upon exact matches to references in the database. Although StrainEst was tested using the same reference database as StrainGE, StrainEst often reported a strain different from the true closest strain in the database or none at all, thus lowering both its precision and recall. However, in these cases, StrainEst still selected a strain with relatively high similarity, as reflected in its high Mash similarities. In contrast, while BIB often picked the closest strain in its database (mean recall 0.94), the selected reference was often a poor proxy given BIB's smaller database, resulting in much lower Mash similarities.

The high performance of StrainGST was especially striking at lower coverages. While StrainGST consistently performed well across all coverages and mixtures, both StrainEst and BIB performed poorly at coverages <1x. StrainGST was the only tool able to recover mixtures of strains present at a 20-fold coverage difference, as reflected by a mean precision of 0.98 and mean recall of 1.0 in the 10x:1x:0.5x benchmark (Supplementary Figure 1). These results highlight the wide dynamic range over which StrainGST was able to correctly identify the closest strains within mixtures, including for the abundance range typically seen for key organisms such as *E. coli* in the human gut.

**C. StrainGR accurately identifies SNVs at low coverages.** StrainGR is unique in its ability to call SNVs across the close reference genomes of strain(s) identified by StrainGST using metagenomic data. Although other tools can identify nucleotide-level differences across sets of samples, they are limited to either marker gene sets, or a single reference, which may be quite distant. To further characterize the ability of StrainGE to call SNVs within low-abundance strains in a metagenomic sample, we introduced random SNVs into sets of *Escherichia* genomes at equal abundance, or unequal abundance. We mixed simulated reads from these strains into a real metagenomic sample, and compared the SNVs called by StrainGR to the known SNVs (Supplemental Figure 2a-b; Supplemental Materials and Methods). StrainGR achieved near perfect precision and recall at identifying true SNVs (>0.95 for coverages 0.5x and above). However, the fraction of the genome where StrainGR was able to make calls (the "% callable genome") was reduced when coverage decreased, or when multiple strains were present, due to there being a greater fraction of shared genome content between references decreasing the unique regions that StrainGE can use for SNV calling. We observed no clear reduction in precision or recall for mixes, either at equal or unequal abundances, highlighting the ability of StrainGR to effectively disentangle SNVs from different strains.

**D. StrainGR accurately identifies large deletions at low coverages.** Because of frequent recombination and horizontal gene transfer in bacteria, patterns of large deletions (gaps) provide an orthogonal line of evidence for strain similarity (10). StrainGR is unique in its ability to call large deletions relative to close reference genomes. In order to benchmark this ability, we introduced random deletions of 5-100kb into *Escherichia* genomes, and mixed simulated reads from these strains into a real metagenomic sample. We then compared the deletions predicted by StrainGR to the known deletions by computing the Jaccard similarity (Supplementary Methods). StrainGR's large deletion predictions closely matched the true deletions, with a Jaccard similarity of approximately 0.8 for coverages 0.5x and higher (Supplementary Figure 2c), and high concordance when examining genome-wide patterns of deletions (example in Supplementary Figure 2d). Multi-mapping reads (due to repeats in the reference genome) reduced the accuracy of calling deletions, as multi-mapping reads that map to a region of the reference that is present, as well as all or part of a deleted region, will not be properly marked as a deletion. When ignoring positions with a majority of multi-mapped reads, the concordance between predicted and true deletions was even higher, reaching a Jaccard similarity score of 0.9 at 10x coverage (Supplementary Figure 2c). The pattern of deletions shared across strains in a dataset should be consistent across all samples to be compared and may reflect evolutionary history, thus providing another key indicator of strain relatedness.

### Supplementary Methods

***StrainGST benchmarking***. We compared the ability of StrainGST to select the closest strain in an *Escherichia* reference database to BIB (4) StrainEst (5). We excluded PathoScope (6) because its database construction process required taxonomy IDs in BLAST's NT database, which have been phased out by NCBI. Sigma was excluded because we could not run the pipeline end-to-end, as we were unable to run steps that depended on MPI for compute parallelism. Where possible, we used the same database to ensure fair comparison. For StrainGST and StrainEst, we used the same 361-strain database. For BIB, since BIB's database construction process did not scale, we generated a smaller database containing 20 representative genomes. To select the 20 representatives, we computed pairwise Mash distances (4) between all 929 genomes used as input into the StrainGST *Escherichia* database and performed hierarchical clustering to obtain 20 clusters. The genome from each cluster with the lowest average distance to all other genomes in its cluster was selected.

We took into consideration StrainEst and BIB calls that reported a strain at >1% abundance relative to other strains in the database, the same threshold used in Albanese and Donati (5). We used pairwise Mash distances to assess how close a reported strain was to the true strain, counting a reported strain as a true positive if it was the closest strain in the database to the strain in the sample. Any other reported strain that was not present in the sample was counted as a false positive. If any of the strains present in the sample were not reported by the tool, it was counted as a false negative.

We ran each tool on 240 spiked metagenomes with 1-4 strains mixed at equal abundance, with average coverage of 0.1x, 0.5x, 1x or 10x; 40 spiked metagenomes with two strains mixed at 10x:1x or 1x:0.1x; and 40 spiked metagenomes with three strains spiked at 10x:1x:0.5x or 1x:0.5x:0.1x. Strains for each sample were randomly selected from NCBI RefSeq and metagenomes were generated as described in the main text Materials & Methods.

***Application of StrainGE using a Salmonella database.*** We constructed a StrainGST database from 877 genomes identified as *Salmonella* in NCBI RefSeq. 177 genomes were retained after database clustering using default settings (clustering genomes with ANI higher than approximately 99.8% ANI to another reference in the database, and keeping a single representative from each cluster). We ran StrainGST with this final database, using default settings.

ANI comparisons between HNK130 and the other members of the Salmonella reference database were approximated using Mash-based k-mer similarity metrics available in StrainGE. BLAST results for HNK130 revealed close hits to *E. coli* rather than *Salmonella*. ANI comparisons between the HNK130 genome and *E. coli* genomes were performed using Chunlab's ANI calculator tool (https://www.ezbiocloud.net/tools/ani) (11).

As a positive control test set to verify that StrainGE works on *Salmonella*, we ran previously published metagenomic datasets where *Salmonella* content was proven (12) against our cleaned *Salmonella* database where the contaminating *E. coli* genomes had been removed.

***Benchmarking of StrainGR SNV calls using simulated data***. To benchmark the ability of StrainGR to call SNVs, we introduced random SNVs into randomly drawn genomes from the NCBI RefSeq complete database, such that the average nucleotide identity to the original reference was 99.9% (approximately 5,000 SNPs). We generated synthetic reads from these genomes and spiked them into a metagenomic sample with no *E. coli* as for other benchmarks (Materials and Methods). We constructed a total of 320 synthetic communities with spiked-in strains at equal abundance, including i) 20 sets for each number of strains per sample (1-4 strains); and ii) 20 sets at each coverage (0.1x, 0.5x, 1x and 10x, corresponding to relative abundances of approximately 0.02x - 1.6x). We also created 20 two-strain communities at 10x:1x and another 20 at 1x:0.5x coverage.

Using these simulated metagenomic samples, we used StrainGR to investigate whether we could correctly identify the synthetically introduced SNPs or deletions, even when mixed within a metagenomic background. For each sample, we prepared a concatenated reference containing the original references used to generate the benchmark sample. We aligned the reads to its concatenated reference, and ran StrainGR to call SNVs. The SNV calls made by StrainGR were compared to the truth (i.e. the known set of mutations introduced into that synthetic strain) using Illumina's som.py (https://github.com/Illumina/hap.py) and each call was classified as either a true positive (TP), false positive (FP), or false negative (FN).

***Benchmarking of StrainGR large deletion predictions using simulated data***

To benchmark the accuracy of StrainGR in predicting large deletions, we created a separate set of 80 synthetic samples based on 20 randomly selected *E. coli* genomes present in the NCBI RefSeq complete database, in which we deleted random blocks of genes (sized 5-100kb), resulting in loss of approximately 7.5% of the total genes in each reference genome. From these synthetic samples, we simulated reads using ART (13) at fixed coverages of 0.1x, 0.5x, 1x and 10x, and mixed the simulated reads with reads subsampled from a real metagenomic sample, as for the SNV benchmarks described above. Reads were aligned to the original reference, and large deletions predicted by StrainGR were compared to true deletions using the Jaccard similarity metric:

$$\text{Jaccard similarity} = \frac{|G_s \cap G_t|}{|G_s \cup G_t|}.$$

$G_s$ is the set of positions in the genome where StrainGR predicted a large deletion, and $G_t$ is the actual, known set of positions for large deletions.
